# RUNX1 haploinsufficiency causes a marked deficiency of megakaryocyte-biased hematopoietic progenitor cells: Mechanistic studies and drug correction

**DOI:** 10.1101/2020.08.23.260281

**Authors:** Brian Estevez, Sara Borst, Danuta Jarocha, Varun Sudunagunta, Michael Gonzalez, James Garifallou, Hakon Hakonarson, Peng Gao, Kai Tan, Paul Liu, Sumedha Bagga, Nicholas Holdreith, Wei Tong, Nancy Speck, Deborah L. French, Paul Gadue, Mortimer Poncz

## Abstract

Patients with familial platelet disorder with a predisposition to myeloid malignancy (FPDMM) harbor germline monoallelic mutations in a key hematopoietic transcription factor RUNX1. Previous studies of FPDMM have focused on megakaryocyte (Mk) differentiation, and platelet production and signaling. However, the effects of RUNX1 haploinsufficiency on hematopoietic progenitor cells (HPCs) and subsequent megakaryopoiesis remains incomplete. To address this issue, we studied induced-pluripotent stem cell (iPSC)-derived HPCs (iHPCs) and Mks (iMks) from both patient-derived lines and a wildtype line modified to be RUNX1 haploinsufficient (RUNX1^+/−^), each compared to their isogenic wildtype control. All RUNX1^+/−^ lines showed decreased iMk yield and depletion of a Mk-biased iHPC subpopulation. To investigate global and local gene expression changes underlying this iHPC shift, single-cell RNA sequencing was performed on sorted FPDMM and control iHPCs. We defined several cell subpopulations in FPDMM Mk-biased iHPCs. Analyses of gene sets upregulated in FPDMM iHPCs indicated enrichment for response to stress, regulation of signal transduction and response to cytokine gene sets. Immunoblotting studies in FPDMM iMks were consistent with these findings, but also identified augmented baseline c-Jun N-terminal kinase (JNK) phosphorylation, known to be activated by transforming growth factor β1 and cellular stressors. J-IN8 and RepSox, small drugs targeting these pathways, corrected quantitative defects in FPDMM iHPC production. These findings were confirmed in adult human CD34^+^-derived stem and progenitor cells transduced with lentiviral *RUNX1* short-hairpin (sh) RNA to mimic RUNX1^+/−^. These mechanistic studies of the defect in megakaryopoiesis in FPDMM suggest druggable pathways for clinical management of thrombocytopenia in affected patients.

**Key points:**

- RUNX1 haploinsufficiency results in a deficiency of megakaryocyte-biased hematopoietic progenitor cells (HPCs).
- RUNX1 haploinsufficiency elevates druggable proinflammatory and TGFβR1-related pathways in HPCs.

## Introduction

The runt-related transcription factor 1 (RUNX1) is critical in definitive hematopoiesis, lymphopoiesis and myeloid differentiation^1–4^. Patients with familial platelet disorder with predisposition to myeloid malignancy (FPDMM) harbor germline monoallelic mutations in *RUNX1* and often present with thrombocytopenia and bleeding^5^. These individuals also have an increased lifetime risk of acquiring myeloid dysplasias and malignancies^6^. Investigations of FPDMM patients have focused on the role of RUNX1 haploinsufficiency (RUNX1^+/−^) in megakaryopoiesis, and platelet number and function^7–18^. The effect of RUNX1^+/−^ on hematopoietic stem and progenitor cell (HSPCs) differentiation has not been well studied though such cells may contribute to both the megakaryocyte (Mk) and platelet defects and oncogenic evolution.

HSPCs sit atop the hematopoietic hierarchy^19–21^, but are not comprised of homogenous populations transitioning through sequential intermediate progenitor stages into all mature blood lineages. Single-cell transcriptomics, single-cell functional assays, and lineage-tracing studies show that phenotypic HSPCs consist of subpopulations with variable long-term self-renewal potential that are transcriptionally and functionally unipotent or biased toward production of certain blood lineages, and undergo early molecular lineage commitment in hematopoiesis^22–26^. In particular, von Willebrand factor-expressing (vWF^+^) hematopoietic stem cells in mice have long-term multilineage reconstitution potential, but are skewed towards the Mk lineage^22,25^. In human hematopoiesis there are Mk-restricted progenitors in the immunophenotypic HSPC and common myeloid progenitor compartments that display in vivo self-renewal and robust platelet-producing activity upon xenotransplantation^27,28^. Mk-biased HSPCs are responsive to stress and inflammation^25,26,29^. The degree and direction of HSPC lineage bias is linked to levels of transcription factor gene modules, such as GATA2/NFE2^24,28^, which together with RUNX1 form key regulatory networks utilized by HSPCs^30,31^.

Heterozygous mutant *Runx1* mice do not recapitulate FPDMM^4,32,33^. Conditional *Runx1* mutant mice display myeloid differentiation defects and paradoxical expansion of HSPCs^3,32,34^. These species differences have focused etiologic studies of FPDMM to RUNX1^+/−^ human induced-pluripotent stem cells (iPSC) that seem to replicate the megakaryocytic defects seen in FPDMM patients^35–37^. Such studies showed that RUNX1^+/−^ results in an ~50% level of final Mks per initial iPSC-derived hematopoietic progenitor cell (iHPC) compared to isogenic controls^36^. A defect in NOTCH signaling has been proposed to be linked to the observed decrease in Mk yield^37^.

We have similarly studied iHPCs and the resultant Mks (iMks) from affected individuals to understand the relationship between the decrease in Mk yield and a defect in HPCs. Our data confirmed the previously observed defect in megakaryopoiesis caused by FPDMM^11,35,36^, but also identified a marked deficiency of a Mk-biased iHPC population. By single-cell RNA sequencing (scRNA-SEQ), this deficiency was associated with an increase in gene sets associated with response to stress, regulation of signal transduction and response to cytokine. We also observed an increase in the stress pathway by upregulation of c-Jun N-terminal kinase (JNK) 2 phosphorylation and increased sensitivity to transforming growth factor beta (TGFβ) 1. Drugs that block these pathways correct the megakaryopoiesis defect. Since iPSCs are embryonic in nature, and may generate an immature blood developmental program, we used an short-hairpin (sh) RNA approach to mimic RUNX1 haploinsufficiency in adult CD34^+^-HSPCs, referred to here as RUNX1 insufficiency (RUNX1^in^), and observed Mk deficiency associated with a decrease in a Mk-biased HSPC population correctable by blocking JNK and TGFβ receptor 1 (TGFβR1) signaling. The mechanistic and therapeutic implications of these findings for patients with FPDMM are discussed.

## Materials and methods

### iPSC lines and mobilized, peripheral-blood-derived, adult CD34^+^ cells

All iPSC constructs studied are noted in Figure Supplement (S) 1. The establishment of line L1 was done with Institutional Review Board approval and carried out in accord with the Helsinki Principles. Granulocyte colony-stimulating factor-mobilized CD34^+^-derived peripheral-blood HSPCs were purchased from the Fred Hutchinson Cancer Research Center Hematopoietic Cell Procurement and Processing Services Core^38,39^.

### Flow cytometry and cell sorting

Flow cytometric analyses were performed on a CytoFLEX LX flow cytometer (Beckman Coulter) and analyzed using FlowJo software v10.6 (Becton-Dickinson). CD235a^+^CD41^+^ surface markers were used to isolate and enumerate iHPCs, whereas CD34^+^CD38^−^CD45RA^−^CD41^+^ surface markers were used to isolate and enumerate adult CD34^+^ HSPCs. Absolute numbers of iPSC- or CD34^+^-derived Mks were determined using annexin V binding in combination with CD41 and CD42 cell surface markers. For a full list of the labeled markers, antibodies and their sources, see Table S1. A FACS Jazz (Becton-Dickinson) was used to sort cells using a 100 μm nozzle. Gating schemes for iHPCs and CD34^+^ HSPCs are provided (Figures S2 and S3).

### Statistical analysis

Statistical analysis was performed using one-way and two-way ANOVA, and data reported as ± 1 standard error of the mean (SEM), using GraphPad Prism version 8 (GraphPad Software). Differential expression analysis of input genes and P-values used to generate Volcano plots were computed in Loupe Cell Browser by negative binomial exact test, which generates P-values that are then adjusted for multiple tests using the Benjamini-Hochberg procedure to decrease the false discovery rate^40^. Differences were considered significant when P values were <0.05.

### Additional methodology

See Supplemental Methods for additional details of RUNX1^+/−^ CD34^+^ HSPC creation and analysis, analysis of hematopoietic differentiation, scRNA-SEQ library preparation, data pre-processing and analysis, and Western blotting and immunofluorescence studies.

### Data Sharing Statement

Data generated during this study have been deposited in the Gene Expression Omnibus (GEO) (GSE149136).

## Results

### Effect of RUNX1^+/−^ on Mk-biased progenitors that emerge with other iHPCs

FPDMM patients display defects in Mks and platelets^11–13,16,17,36^. Whether these phenotypes stem from defects in earlier HSPC populations is unclear. HSPCs displaying Mk-biased lineage commitment have been reported in mice and humans^22,25,27,28^. Whether these early progenitors are affected in RUNX1^+/−^ has yet to be examined. To investigate the effect of RUNX1^+/−^ on early Mk-biased HPCs in FPDMM patients, we utilized three distinct RUNX1^+/−^ iPSC lines and their isogenic control (Figure S1). L1 was from an FPDMM patient harboring a previously characterized monoallelic splice-site acceptor mutation in *RUNX1* intron 3^41^, and its isogenic mutation-corrected line, L1-C. A second set included a previously-established wildtype line (WT) altered to introduce this splice mutation (WT-L1). A third set included RUNX1^+/−^ L2 with a C→A transversion in *RUNX1* exon 7 that introduced a stop codon in RUNX1’s transactivation domain^*35*^ and its isogenic control L2-C^35^. These six lines were exposed to a directed hematopoietic differentiation program to generate iHPCs, which arise as free-floating CD235^+^CD41^+^ cells on Day 7 of culture (Figure 1A and ^42,43^) as these markers enrich for multipotent progenitor cells^43,44^. They were then differentiated into Mks (Figure 1A)^45^. All three RUNX1^+/−^ lines showed decreased iMk yield per iHPC (Figure S4A). Reduction in RUNX1 levels in terminal iMks was validated for lines L1 and L2 (Figure S4B), consistent with FPDMM patients^11^ and prior iMk studies^35–37^.

**Figure 1.**
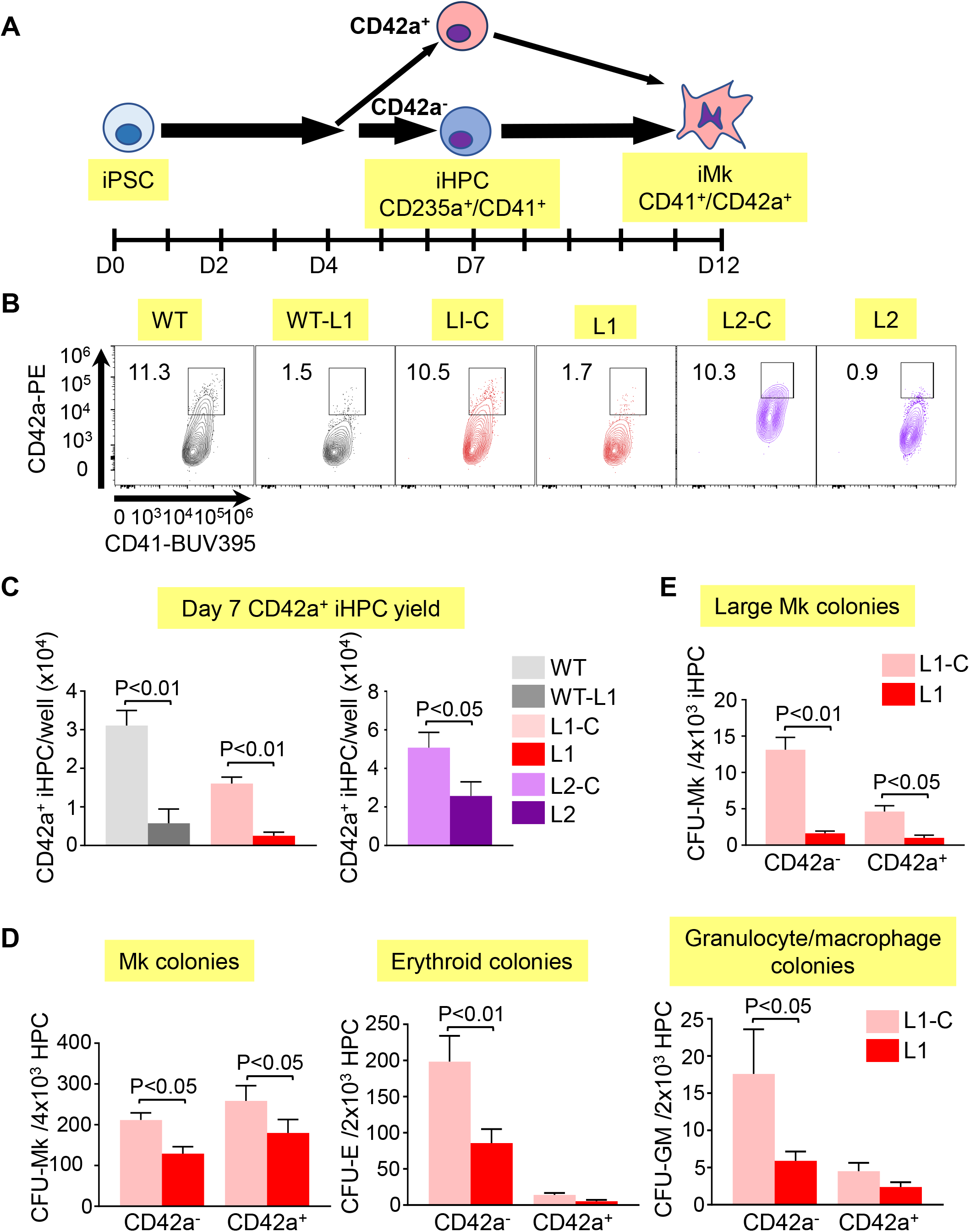
Deficiency in a Mk-biased CD42a^+^ subpopulation in RUNX1^+/−^ iHPCs. (A) Schematic of megakaryopoiesis in a human iPSC system highlighting generation of iHPCs and iMks. On Day 7 after exposure of adherent iPSCs to a directed differentiation protocol, free-floating multipotent CD235^+^CD41^+^ iHPCs emerge that include CD42a^−^ and CD42a^+^ iHPC subpopulations. (B) Flow cytometric analysis of Day 7 iHPCs showing the CD42a^+^ iHPC subpopulation that is present at similar frequencies in all control lines, but is severely reduced in all RUNX1^+/−^ lines. (C) is quantitation of (B) showing the mean number of CD42^+^ iHPCs detected normalized per well of iPSCs seeded ± 1 standard error of the mean (SEM). N = 5-15 studies/arm. P value was determined by Student’s t-test. (D) Colony-forming assays of L1-C versus L1 iHPCs, showing quantitation of mean numbers of Mk, erythroid, and granulocyte/macrophage colonies normalized to the number of Day 7 iHPCs seeded ± 1 SEM. N = 4 studies/arm. P value was determined by Student’s t-test. (E) Same as (D), but for large Mk colonies (>20 Mks/colony).

We tested whether we could detect a Mk-biased iHPC population, analogous to the adult human Mk-biased HSPC population,^27,28,46^ and whether this population is deficient in RUNX1^+/−^. We evaluated iHPCs for intracellular stores of vWF, a known Mk- and endothelial cell-associated marker, shown to be enriched in murine Mk-biased HSPCs^22^. Using intracellular flow cytometry and immunofluorescence (IF) microscopy, we detected vWF expression in all normal and RUNX1^+/−^ iHPCs with no significant difference (Figures S5A and S5B). Similarly, although we detected expression of Mk-associated α-granule protein TREM-like transcript-1 (TLT-1) in iHPCs, we did not see any significant subpopulation of iHPCs based on TLT1 nor a difference between the controls and RUNX1^+/−^ iHPCs (Figure S5C, and not shown).

We did identify a novel CD235^+^CD41^+^CD42a^+^ iHPC subpopulation (Figures 1A to 1C, and Figures S3A and S3B). CD42a is the Mk-specific protein, glycoprotein (GP) IX, a component of the Mk-specific GPIb/IX receptor^47,48^. We sorted CD42a^−^ and CD42a^+^ subpopulations from CD41+CD235+ iHPCs and seeded equal numbers in Mk-, erythroid and myeloid-colony forming assays. Both control and RUNX1^+/−^ CD42a^−^ iHPCs, displayed trilineage potential, while both CD42a^+^ iHPCs produced near-exclusive Mk colonies (Figure 1D). For all RUNX1^+/−^ iPSC lines, there was a 50-80% decrease in CD42a^+^ iHPCs compared to control (Figures 1B and 1C, and Table S2). In addition, RUNX1^+/−^ iHPCs had a near-absence of large Mk colony formation (Figure 1E), suggestive of a proliferative defect in these cells. Whether this defect delayed CD42a^+^ iHPC emergence from RUNX1^+/−^ iPSCs was tested and showed defective CD42^+^ iHPCs continuing beyond Day 7 (Figure S6). These results suggest RUNX1^+/−^ severely affects the development of Mk-biased progenitors emerging concurrent with other iHPCs.

### scRNA SEQ identifies novel deregulated pathways in RUNX1^+/−^ iHPCs

To gain insights into the global and local transcriptomic changes underlying defective production of CD42a^+^ iHPCs in RUNX1^+/−^, we performed scRNA SEQ on sorted CD42a^−^ and CD42a^+^ iHPCs from L1 and L1-C. There were on average 11,328 reads per cell with an average of 2,081 genes expressed. To identify and analyze distinct subpopulations in the control and RUNX1^+/−^ iHPCs, dimension reduction was performed and cell clustering was visualized using uniform manifold approximation and projection (UMAP)^49^. Based on differentially expressed (DE) genes, 8 transcriptomic cell clusters were defined in CD42a^+^ iHPCs in both control and RUNX1^+/−^ iHPCs (Figure 2A and Table S3). To identify changes due to RUNX1^+/−^, we compared the numbers of DE genes between clusters. L1-C and L1 CD42a^+^ iHPCs, clusters 1, 3, 4 and 7 showed the highest numbers of DE genes (Figure S7A). Functional enrichment profiling of genes upregulated in L1-C iHPCs clusters^50^ was consistent with previous scRNA-SEQ studies showing a large contribution of cell cycle genes to transcriptional heterogeneity observed in differentiating hematopoietic cells^51^, particularly stem and progenitor cells^52,53^. L1-C CD42a^+^ iHPCs, clusters 1 and 3 were enriched in cell cycle-related biological processes, whereas cluster 7 showed broad enrichment in myeloid- and especially Mk/platelet-associated genes. Cluster 4 in L1-C CD42^+^ iHPCs was enriched for both types of biological processes (Figure 2B), suggesting an intermediate state of maturation.

**Figure 2.**
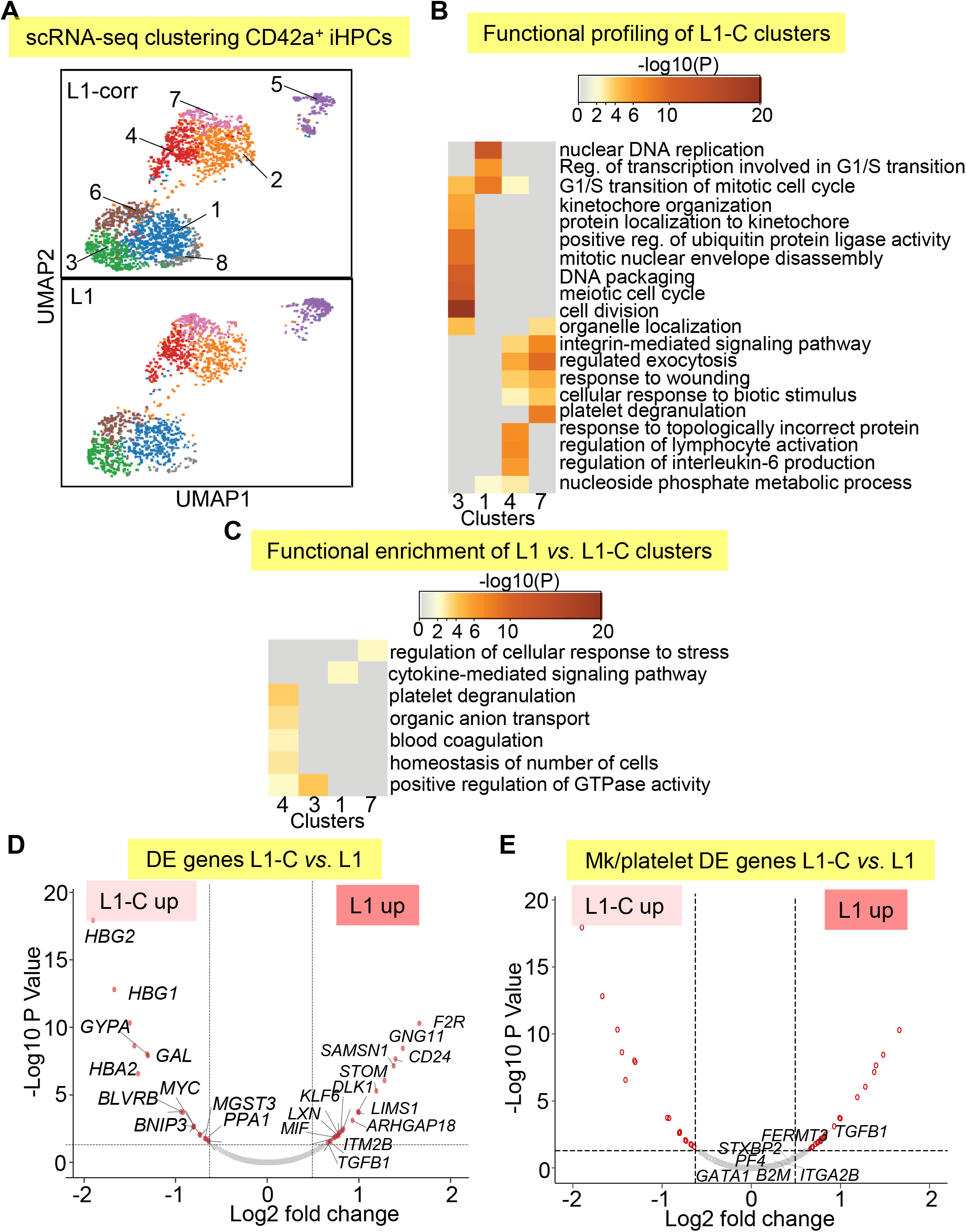
scRNA-SEQ analysis of Mk-biased CD42^+^ iHPCs. (A) Identification and visualization of transcriptional heterogeneity within sorted L1-C and L1 CD42^+^ iHPCs by dimensionality reduction using UMAP. (B) and (C) Gene ontology analyses using lists of genes from select clusters with highest detected numbers of DE genes to show functional enrichment in (B) control CD42a^+^ iHPC clusters only or (C) processes upregulated in L1-C versus L1 CD42^+^ iHPCs. (D) and (E) Volcano plots showing significantly changed genes (D) among all DE genes or (E) focusing on Mk/platelet-associated genes when comparing L1-C to L1 CD42a^+^ iHPCs across all clusters. To improve clarity, only select genes are shown and labeled (see Tables S5 and S6 for a full list). Filled red circles denote significantly changed genes, whereas unfilled red circles indicate DE genes that are not Mk/platelet-associated genes.

Most genes from a curated list of 133 known Mk-associated genes (Table S4) did not show significant differences in global DE analysis between CD42a^−^ and CD42a^+^ iHPCs subpopulations (Figures S7C, S8 and S9A, and Table S5). We believe that clusters enriched in cell cycle genes are proliferating iHPCs, whereas those deficient are likely to be iHPCs further along in maturation (Figures S10 and S11, and Table S6). In support, gene-set enrichment analysis (GSEA) indicates that in both lines, CD42a^+^ iHPCs show enrichment of Mk progenitor- and myeloid-associated genes, and depletion of erythroid-associated genes, consistent with an early Mk-bias (Figure S12). Analysis of gene sets upregulated in L1 CD42a^+^ iHPC clusters revealed enrichment of several processes, including platelet degranulation, cytokine-mediated signaling, and regulation of cellular response to stress (Figure 2C and Table S7). Many genes found in these enriched processes were also DE in global analyses when comparing all cell clusters from L1-C and L1 CD42^+^ iHPCs (Figures 2D and 2E, and Table S8).

For CD42a^−^ iHPCs, RNA SEQ was done on N=2,909 L1-C cells and 2,270 L1 cells. UMAP clustering defined 6 transcriptomically distinct cell clusters (Figure 3A). Functional profiling of control L1-C CD42a^−^ iHPC clusters showed a similar pattern of distinct cluster-specific enrichment of cell cycle and erythroid, myeloid and Mk/Plt gene sets (Figure 3B). Interestingly, DE genes in RUNX1^+/−^ CD42a^−^ iHPC clusters showed broad enrichment for stress-associated processes (Figure 3C), although L1 CD42^+^ iHPCs showed a greater number of DE and upregulated genes than L1 CD42a^−^ iHPCs (Figures 3D and 3E, Figure S7A-B), wherein functional profiling revealed a similar enrichment for stress-associated and pro-inflammatory processes (Figure S9B).

**Figure 3.**
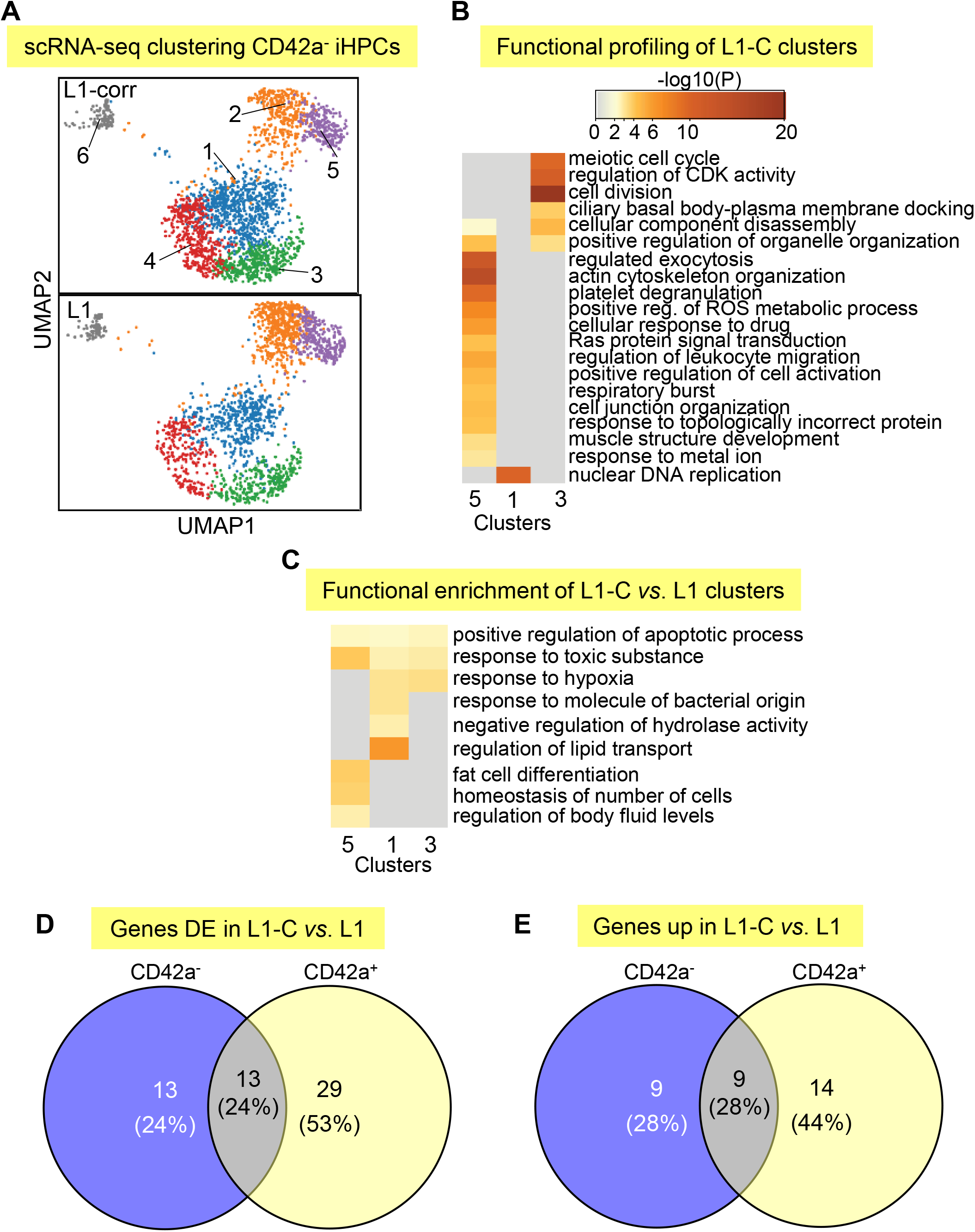
scRNA-SEQ analysis in CD42a^−^ iHPCs and comparison of deregulated genes in RUNX1^+/−^ CD42a^−^ and CD42a^+^ iHPCs. (A) Identification and visualization of transcriptional heterogeneity within sorted L1-C and L1 CD42^−^ iHPCs by UMAP. (B) and (C) Gene ontology analyses using lists of genes from select clusters with highest detected numbers of DE genes to show functional enrichment in (B) control CD42a^−^ iHPC clusters only or (C) processes upregulated in RUNX1^+/−^ L1 compared to L1-C CD42a^−^ iHPCs. (D) and (E) Venn diagrams comparing similarities and differences in DE and upregulated genes in RUNX1^+/−^ CD42^−^ and CD42^+^ iHPCs for (D) all DE genes or only (E) upregulated genes.

Our analysis of Mk/platelet-associated genes in RUNX1^+/−^ CD42^+^ iHPCs indicated that *TGFB1* was among the highest DE genes overall (Figure 2E) and that thrombospondin 1 (*THBS1*), a negative regulator of megakaryopoiesis known to be involved in regulating TGFβ1 activity^54,55^, was also among the highest DE genes in cluster-specific analyses (Figure S13). Together, local and global analyses of genes in RUNX1^+/−^ L1 iHPCs revealed upregulated TGFβ1-related processes.

### Augmented TGFβ1 and JNK signaling in RUNX1^+/−^

Many of the augmented cytokine and cellular stress response genes in the RUNX1^+/−^ CD42a^+^ iHPCs involve TGFβR1-associated pathways^56–61^. Therefore, their deficiency could be due to elevated TGFβ1 and/or TGFβR1-associated levels or signaling. As TGFβ1 is released in the marrow predominantly by Mks^62,63^, we tested whether RUNX1^+/−^ iMks release excess TGFβ1; however, instead of elevated levels, we found decreased TGFβ1 levels in RUNX1^+/−^ iMks (Figure 4A). With correction of quantitative defects in iMk yield in RUNX1^+/−^ iMks cultures, TGFβ1 yield per iMk was similar in L1 compared to L1-C iMk cultures (Figure 4B). We next tested whether defective RUNX1^+/−^ CD42^+^ iHPC production and iMk yield were due, in part, to enhanced sensitivity to TGFβ1, isogenic control and RUNX1^+/−^ iHPC cultures were exposed to increasing doses of recombinant human (rh) TGFβ1 protein, which is known to inhibit Mk proliferation^64^. RUNX1^+/−^ cultures showed greater sensitivity to rhTGFβ1 (Figure 4C), which could be rescued by treatment with anti-TGFβ1 blocking antibodies^65^ (Figure 4D).

**Figure 4.**
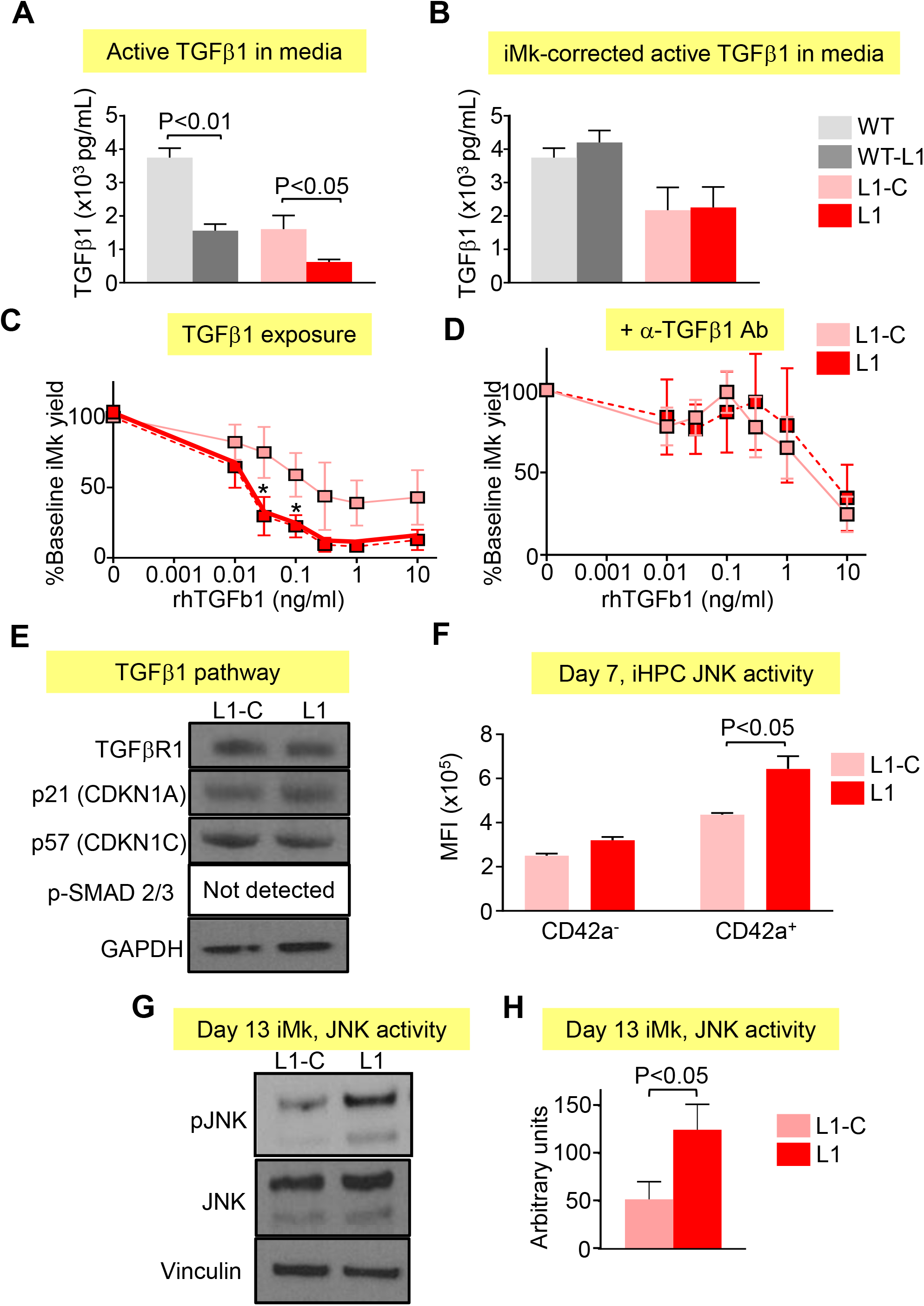
Augmented TGFβ1 and JNK signaling in RUNX1^+/−^ iHPCs and iMks. (A) and (B) ELISA of TGFβ1 levels in iMk culture conditioned medium. Cell culture supernatants were collected from L1-C and L1 Day 13 iMk liquid cultures and used to measure levels of active TGFβ1. (A) Mean ± 1 SEM of TGFβ1 without normalization for differences in iMk number and (B) with normalization for iMk numbers. N = 5 studies/arm with P values determined by Student’s t-test. (C) Effect of TGFβ1 exposure on Mk yield from L1-C and L1 iHPCs. Cultures were exposed to rhTGFβ1 for 5 days at indicated doses, and Mk yield was quantified per input iHPC as in Figure 1C. Results are shown as a percent of untreated iMk yield. (D) Effect of 20 μg/mL anti (α) -TGFβ blocking antibodies (Abs) on Mk yield suppression resulting from TGFβ1 exposure. Mean ± 1 SEM. N = 6 studies/arm with P values calculated two-way ANOVA. (E)-(G) are immunoblots. (E) Day 13 iMk proteins with antibodies directed against TGFβR1, p21, p57 and pSMAD2/3 with GAPDH as a loading control. (F) Day 7 iHPC immunostained with antibodies against phosphorylated (p) JNK. Mean ± 1 SEM. N = 4 studies/arm with P values calculated Student’s t-test. (G) Elevated JNK phosphorylation in Day 13 L1 iMks. These iMks were solubilized and immunoblotted with antibodies against pJNK, total JNK and vinculin as a loading control. (H) Experiments in (G) were scanned and quantified optical density. Mean ± 1 SEM is shown with N = 6 studies/arm with P values calculated by Student’s t-test.

We examined which components of TGFβ1 signaling pathway were responsible for enhanced TGFβ1 sensitivity in RUNX1^+/−^ cells by immunoblot studies on Day 13 iMks, focusing on canonical TGFβ1 signaling pathways (Figure 4E). We did not detect differences in levels of TGFβR1, p21 and p57. SMAD 2/3 protein or phosphorylation levels were not detected in iMks from both L1-C and L1 by immunoblotting and by intracellular flow cytometry of iHPCs (Figure S14). Therefore, we next evaluated phosphorylation of c-Jun N-terminal kinase (JNK), a stress responsive mitogen-activated protein kinase (MAPK) that is known to be activated by TGFβ1^66^. We identified a significant elevation of JNK phosphorylation in RUNX1^+/−^ CD42^+^ iHPCs (Figure 4F) and iMks (Figures 4G and 4H) not due to JNK protein levels (Figure 4G).

### Drug intervention to correct iMK deficiency from RUNX1^+/−^ iHPCs

To determine whether decreasing JNK signaling in differentiating RUNX1^+/−^ iHPCs would correct defective megakaryopoiesis, we evaluated the effect of JNK inhibition on iMk using the small molecule JNK inhibitors, J-IN8^67^ and J-IX^68^. Treatment of L1 iHPCs with J-IN8 improved iMk yield, although J-IX did not (Figures 5A and 5B), which may be related to the relatively reduced potency and increased reversibility of J-IX inhibition^68^.

**Figure 5.**
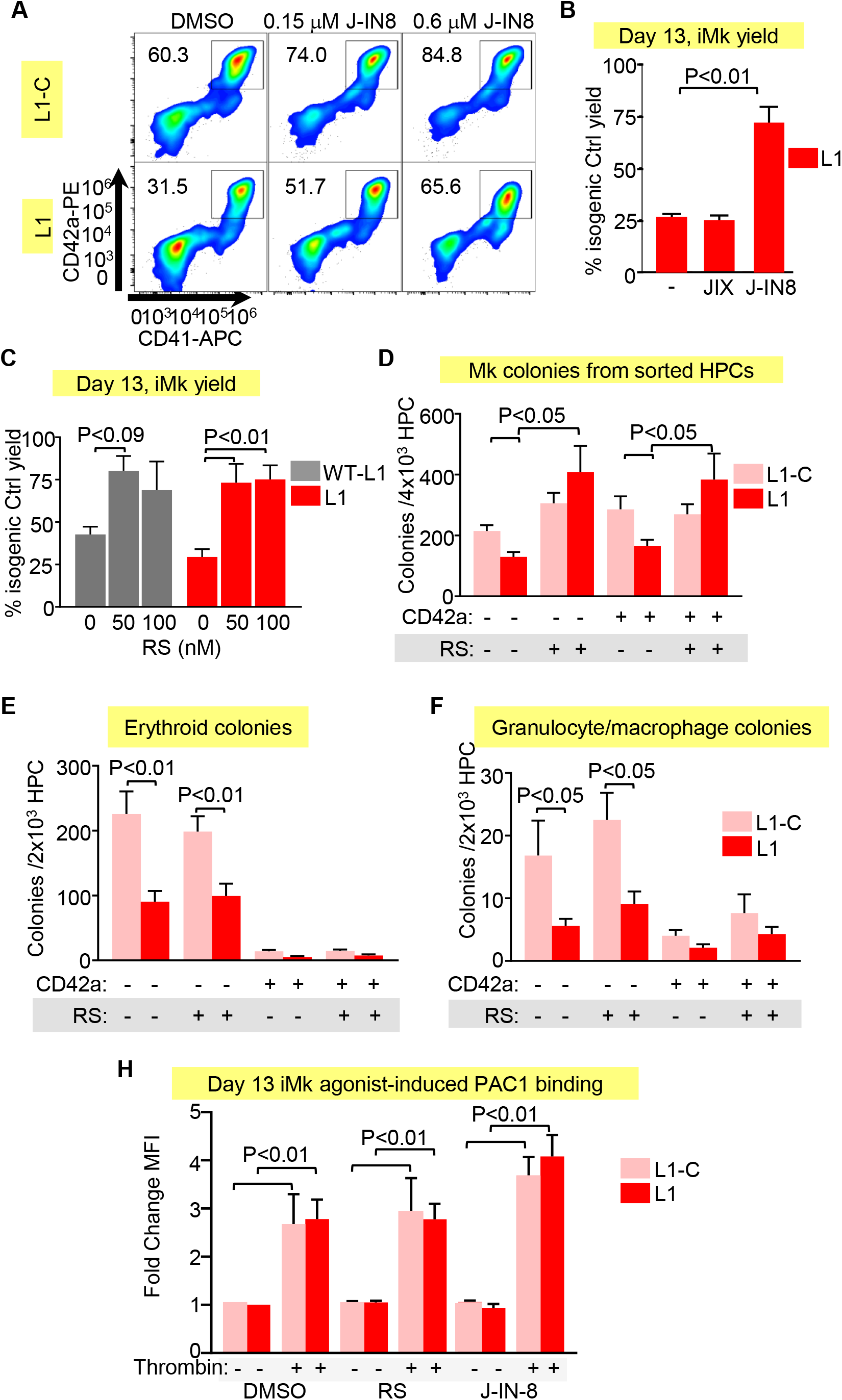
Drug intervention to correct iMK deficiency from RUNX1^+/−^ iHPCs. (A) and (B) Representative flow cytometric analysis L1-C and L1 Day 13 iMks cultures treated with DMSO or J-IN8 and then stained with antibodies against Mk surface markers CD42a and CD41. (B) Mean ± 1 SEM quantitation of the effect of 0.6 μM JNK inhibitors (J-IN8 or JIX) on iMk yield as % DMSO-treated L1-C controls. N = 4 studies/arm with P values calculated one-way ANOVA. (C) Quantitation of the effect of 0.1 μM RS on iMk yield as in (B). N = 4-6 studies/arm with P values calculated one-way ANOVA. (D)-(F) Colony-forming assays performed using sorted L1-C and L1 CD42a^−^ and CD42a^+^ iHPCs treated with DMSO or RS (0.1 μM). Sorted cells were seeded in megacult media for (D) Mk colonies, or methacult media for (E) erythroid colonies, and (F) granulocyte/macrophage colonies. Mean ± 1 SEM is shown with N = 4 studies/arm with P values calculated two-way ANOVA. (H) Quantitation of response of agonist-induced PAC-1 binding in drug-treated L1-C and L1 iMks. After 5 days of culture in the presence DMSO, RS or J-IN8, Day 13 iMks were collected, stained with Mk surface markers, and an integrin α_IIb_β_3_ activation-dependent monoclonal antibody PAC-1, and stimulated with 0.1U/mL thrombin as described^45^. Mean ± 1 SEM is shown with N = 3 studies/arm with P values calculated two-way ANOVA.

We next tested whether targeting TGFβ1 signaling would improve iMk yield from RUNX1^+/−^ iHPCs. RepSox (RS), a small compound TGFβR1 inhibitor, promotes human megakaryopoiesis and increases RUNX1 expression in vitro and in vivo-platelet release in mice^69^. RS-treatment improved iMk yield from L1^−^ iHPCs to near L1-C levels (Figure 5C). Further, in colony-forming assays on sorted CD42^−^ and CD42^+^ iHPCs, RS increased the number of iMk colonies, but did not significantly affect the numbers of erythroid and myeloid colonies (Figures 5D-F). We also tested the effect of these compounds on agonist-induced integrin α_IIb_β_3_ activation as determined by PAC-1 binding on Day 13 iMks. Thrombin-stimulated Day 13 iMks cultured with J-IN8 or RS showed no suppression of PAC-1 binding compared to dimethyl sulfoxide (DMSO) vehicle-only controls, suggesting these drugs improve differentiation, but do not interfere with agonist-induced integrin activation signaling (Figure 5H).

### Studies of RUNX1^in^ adult CD34^+^ HSPCs

FPDMM platelet defect and oncogenic risk involve adult hematopoietic tissues^70^. To determine whether our findings in iHPCs are applicable to adult hematopoiesis, we transduced mobilized, peripheral-blood CD34^+^ HSPCs with lentiviruses expressing *RUNX1*-targeting shRNAs (Figure S15). Both *RUNX1* shRNAs 386 and 813 lentiviruses reduced by ~50% both *RUNX1* message (Figure 6A) and Mk yield in liquid culture conditions (Figure 6B) compared to non-targeting (NT) shRNA controls.

**Figure 6.**
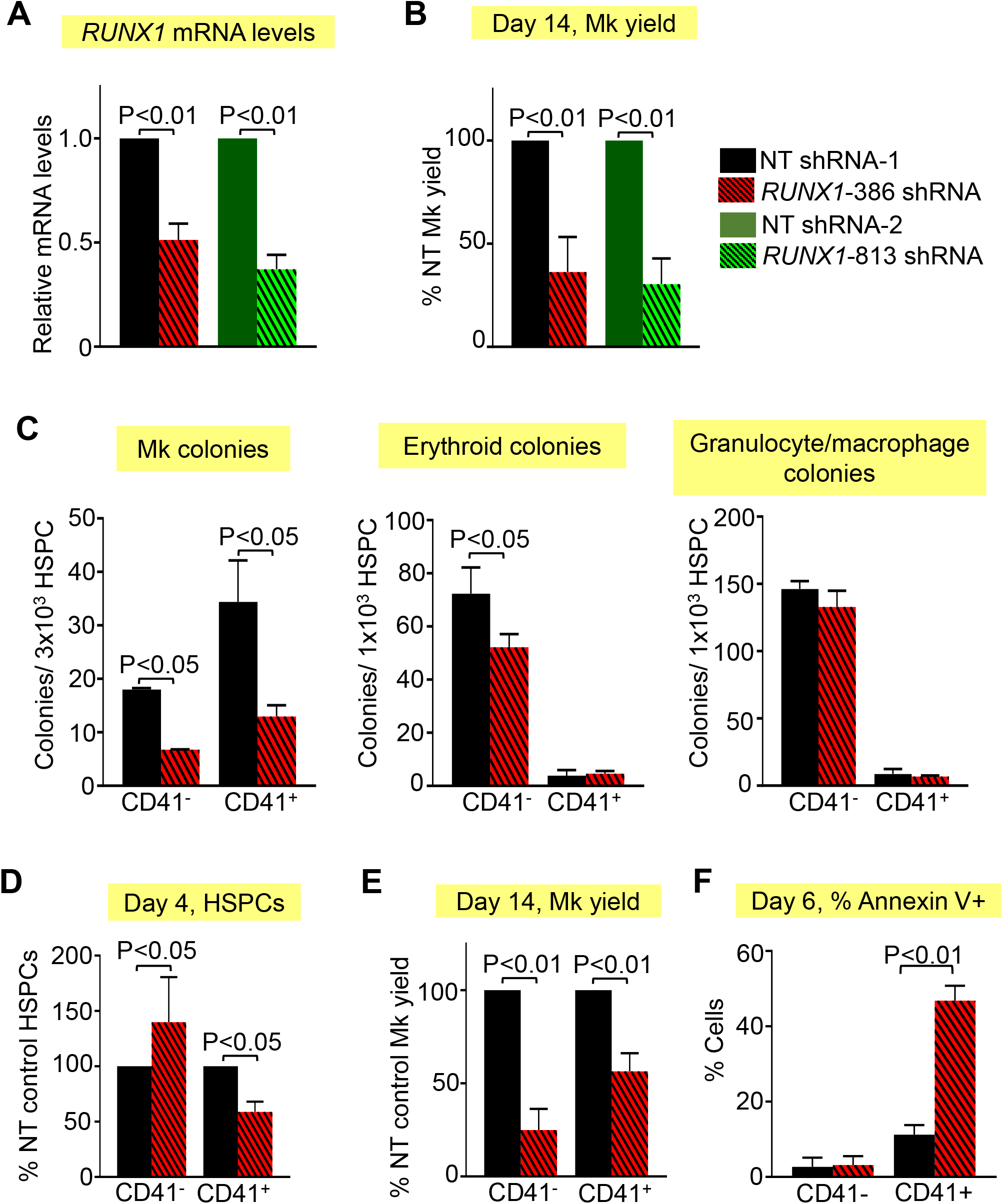
Studies of RUNX1^in^ adult CD34^+^ HSPCs. *RUNX1*-targeting or NT shRNA lentiviral (Figure S2) transduced CD34^+^ HSCs. (A-F) Mean ± 1 SEM with P values calculated by Student’s t-test. (A) Quantitation of *RUNX1* mRNA levels in Day 14 Mk cultures by quantitative PCR with *RUNX1* levels in NT control shRNA-treated cells set as 1. N = 3-5 studies/arm. (B) Day 14 Mk yield ± 1 SEM from *RUNX1* shRNA-transduced CD34^+^ HSPC cultures as a percentage of NT control shRNA. N = 3-5 studies/arm. (C) Colony-forming assays performed using sorted NT- and *RUNX1* shRNA-expressing CD41^−^ and CD41^+^ HPSCs. Sorted cells were seeded in megacult media for Mk colonies or methacult media for erythroid colonies, and granulocyte/macrophage colonies. Mean ± 1 SEM are shown for N = 4 studies/arm. (D) Quantitation of the number of Day 4 CD41^−^ and CD41^+^ HSPCs in NT and *RUNX1*-shRNA lentiviral transduced cells as a percentage of NT shRNA control HSPCs. N = 3-5 studies/arm. (E) Mean ± 1 SEM of Day 14 Mks from sorted CD41^−^ and CD41^+^ HPSCs as a percentage of NT control. N = 4 studies/arm. (F) Mean ± 1 SEM of percent Annexin V positive cells in Day 6 sorted CD41^−^ and CD41^+^ HSPCs.

To identify an immunophenotypic Mk-biased HSPC subpopulation from the overall CD34^+^ HSPC population, we sorted for CD34^+^CD38^−^CD45RA^−^CD41^+^ HSPCs (CD41^+^ HSPCs) and CD34^+^CD38^−^ CD45RA^−^CD41^−^ HSPCs (CD41^−^ HSPCs). We demonstrate that CD41^+^ HSPCs almost exclusively form Mk colonies, whereas CD41^−^ HSPCs displayed multilineage potential (Figure 6C). On Day 4 of culture, total numbers of *RUNX1*-386 shRNA-transduced CD41^+^ HSPCs were reduced and CD41^−^ HSPCs increased compared to non-targeting shRNA controls (Figure 6D). When we seeded equals numbers of sorted cells, RUNX1^in^ introduced by shRNA-transduction resulted in reduced Mk yield both in the CD41^−^ and CD41^+^ HSPCs (Figure 6E). Thus, both Mk-biased and not biased HSPCs contribute to the observed decrease in megakaryopoiesis. *RUNX1*-386 shRNA also significantly reduced the Mk colony number, but erythroid and myeloid yields were either minimally affected or unaffected (Figure 6C). This differs from the pan reduction in terminal hematopoietic lineages seen using RUNX1^+/−^ iHPCs (Figure 2D), but is similar to what is observed in FPDMM patients where cytopenia is limited to platelets^71^. Further analysis of sorted CD41− and CD41+ HSPCs on Day 6 of culture showed a marked enhancement of apoptotic cells in RUNX1^in^ CD41+ HSPCs (Figure 6F).

We next determined whether small molecule inhibitors of JNK and TGFβR1 signaling could promote Mk yield from RUNX1^in^ HSPCs introduced by transduction with lentiviral shRNA. Sorted, mobilized peripheral blood CD34^+^ HSPCs were transduced with NT or *RUNX1*-targeting lentiviral shRNA and showed that both J-IN8 and RS corrected Mk yield per initial HSPC, but that J-IN8 displayed no statistically significant additive effect with RS on Mk yield (Figure 7A and data not shown). RS corrected the yield of Mks from both RUNX1^in^ CD41^−^ and CD41^+^ HPCs (Figure 7B), but did not affect erythroid or myeloid cells (Figures 7C and 7D, respectively). Moreover, a different TGFβR1 inhibitor, Galunisertib (GS), in clinical development^72^, also corrected Mk yield (Figure 7A).

**Figure 7.**
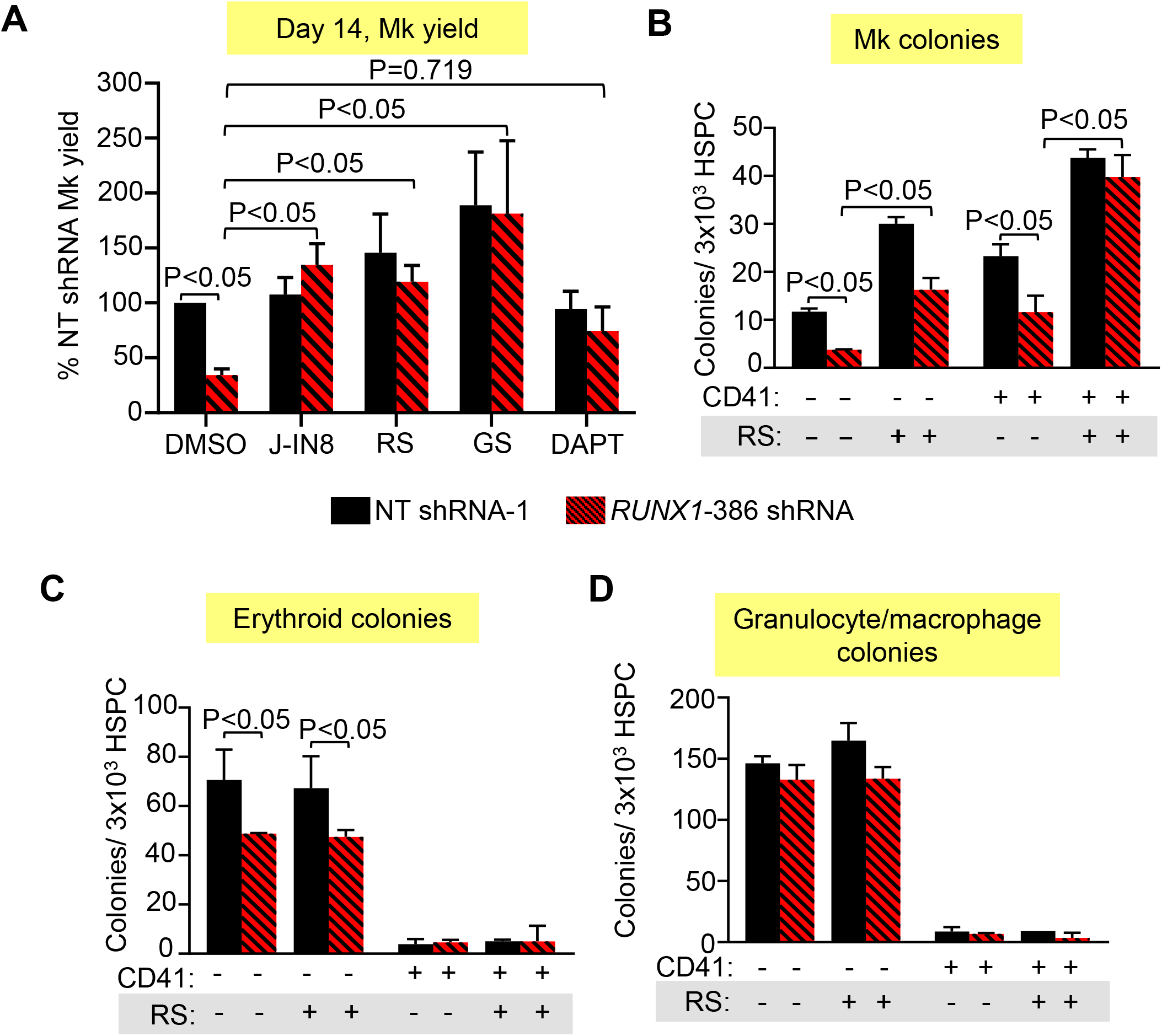
Effect of drugs on megakaryopoiesis in RUNX1^in^ adult CD34^+^ HSPCs. Studies as in Figure 6, but with added J-IN8, RS, GS, and DAPT. In (A) through (D), mean ± 1 SEM are shown. N = 4 studies/arm. P values were calculated by two-way ANOVA. (A) Drug effects on correcting the Mk yield per HSPC of *RUNX1*-versus NT-shRNA lentiviral-transduced CD34^+^ HSPCs after exposure to J-IN8 (0.1 μM), RS (0.1 μM), GS (0.1 μM) or DAPT (10 μM) added Days 5-14. (B-D) Effect of RS on colony-forming assays performed using sorted NT- and *RUNX1* shRNA-expressing CD41^−^ and CD41^+^ HPSCs. Sorted cells were seeded in megacult media for (B) Mk colonies, or methacult media for (C) erythroid colonies, and (D) granulocyte/macrophage colonies.

### Previously defined defects in RUNX1^+/−^ Mks

Most groups studying RUNX1^+/−^ have focused on the observed final Mk yield from iHPCs^36,73^ and from CD34^+^ HSPCs^11,18^. Our group and others have defined defects in multiple late cytoskeletal proteins (e.g., non-muscle myosins IIA and IIB, and myosin light chain 9) ^36^; granule contents or granule trafficking proteins (e.g., PF4, PLDN, RAB1B)^15,17^; platelet activation signaling proteins (e.g., ALOX12 or MYL9)^10^ or Mk-lineage-enriched transcription factors (e.g., NFE2)^13^. A recent study using L2 iPSCs showed that NOTCH4 downregulation in iMks by RUNX1 was defective in RUNX1^+/−^, and that targeting NOTCH signaling with γ-secretase inhibitor DAPT corrected defective megakaryopoiesis^37^. We confirmed that RUNX1 regulates PF4 levels using lentiviral shRNA transduction in adult CD34^+^ HSPCs to introduce RUNX1^in^ (Figure S16A). However, we did not detect elevation of NOTCH4 in RUNX1^+/−^ iMks, studying line L1 (Figure S16B). We also tested the effect of γ-secretase inhibitor DAPT in adult CD34^+^ HSPCs with *RUNX1* shRNA, and saw no significant improvement in Mk yield (Figure 7A). The basis for this difference in NOTCH4 role in RUNX1^+/−^ iMk defect is unclear, but may be related to the use of different RUNX1^+/−^ iPSC lines in the two studies.

## Discussion

FPDMM patients are at high risk for oncogenic progression, but initially manifest predominantly quantitative and qualitative platelet defects. Thus, many investigators have focused on the role of RUNX1 in Mks and platelets. How these insights in platelets may translate into other aspects of the disease is unclear. Recent reports have shown that the Mk lineage emerges, in part, early at the HSPC level, and that a subset of these cells display Mk features and a bias towards terminal Mks^21^. We reasoned that RUNX1 activity in HSPCs promotes Mk-biased differentiation and that targeting effected pathways elevated in these cells might correct at least the Mk and platelet defects. To this end, we explored several approaches, including an FPDMM iPSC-derived model of Mk-biased differentiation and shRNA-derived RUNX1^in^ in adult CD34^+^ HSPCs. Both studies support that RUNX1^+/−^ results in a deficiency of Mk-biased progenitors which contribute to the observed Mk defect and that these Mk-biased progenitor defects are associated with elevations in druggable stress and proinflammatory pathways.

Prior studies of single cell transcriptomics, lineage tracing and in vivo xenotransplantation assays provided evidence that the Mk lineage can be specified at the level of HSPCs^19,21^, and that differentiation directly toward the Mk/platelet lineage occurs under steady-state conditions and in response to inflammatory stimuli^25,29^. In mice, VWF expression marks adult HSCs with Mk-biased differentiation^22^. In humans, we detected VWF expression in both Mk-biased and -unbiased iHPCs, and both were unaffected by RUNX1^+/−^. Since CD41 and CD42 surface markers were reported to enrich for human common myeloid progenitors (CMPs) with Mk-bias^27^, and our iPSC system produces functionally multilineage primitive CMPs^43,44^, we examined whether CD42 surface expression can enrich for Mk-biased CD235^+^CD41^+^ HPCs and found that CD42^+^ Mk-biased iHPCs emerge concurrently with multipotent iHPCs. Our single cell transcriptomic data revealed the heterogenous nature of CD42^+^ iHPCs along with enrichment for several Mk-associated genes and depletion of cell cycle-associated genes. Comparison of our control iHPC transcriptomic data with published gene sets of terminal Mks^74,75^, Plts^75^, and Mk progenitors (MkPs)^76,77^, monocyte dendritic cell progenitors (MD-P), granulocytes (Gran), and erythrocyte progenitors (EryP)^77^ suggests that although most known Mk-associated genes are not significantly upregulated, CD42^+^ iHPCs are enriched for MkP- and granulocyte-associated genes, further supporting that these cells are not just early maturing Mks (Figure S12 and Table S5).

We also investigated Mk bias using adult CD34^+^ HSPCs mobilized into the peripheral blood. We found that a subpopulation of immunophenotypic CD34^+^CD38^−^ HSPCs express CD41 and are Mk-biased in colony assays. The frequency of adult CD34^+^CD38^−^CD41^+^ progenitor cells (~25%) reported here is higher than prior reports in freshly isolated bone-marrow HSPCs^27,78^, likely due to longer time in HSPC expansion culture media containing fms-like tyrosine kinase 3 ligand (FLT3L), thrombopoietin (TPO) and stem cell factor (SCF), which also promotes Mk differentiation. The lentiviral shRNA approach utilized here achieved ~50-70% reduction of RUNX1 levels in terminal adult Mks, and a Mk yield defect as anticipated. We show that RUNX1^+/−^ causes marked defects in production of iMk progenitors; however, Mk-bias is intact in colony assays in RUNX1^+/−^ CD42^+^ iHPCs, similar to control CD42^+^ iHPCs, suggesting RUNX1 promotes early iHPC to iMk commitment, but is not required for iMk bias itself.

scRNA-SEQ revealed that RUNX1^+/−^ CD42^+^ iHPCs display transcriptional deregulation associated with stress, immune and cytokine response pathways. Among DE Mk-associated genes we identified upregulation of *TGFB1* and THBS1, both known for playing a role in TGFβR1 signaling^54,55^, suggesting elevated TGFβ1 signaling contributes to the underlying defect in RUNX1^+/−^ iHPCs. Other upregulated genes in RUNX1^+/−^ iHPCs, such as *KLF6*, *TIMP3*, and *HIPK2* are reportedly associated with or induced by TGFβ1 signaling^56,57,60^. While we saw no elevation of TGFβ1 levels in conditioned media, RUNX1^+/−^ iHPCs displayed enhanced sensitivity to TGFβ1 and enhanced baseline JNK2 phosphorylation, known to be activated by TGFβ1^66^. Further investigation is required to determine how RUNX1 regulates these genes, the role of these molecules in the CD42^+^ iHPC defect, and whether these pathways offer druggable targets in FPDMM patients that would improve the platelet defect and perhaps the oncogenic proclivity. We treated RUNX1^+/−^ HPCs with TGFβR1 and JNK inhibitors to determine if TGFβ1-associated signaling played a role in our embryonic model of FPDMM, and found a significant correction of megakaryopoiesis. Additionally, treatment of RUNX1^+/−^ HPCs with a TGFβR1 inhibitor in colony assays showed selective improvement of the Mk progenitor defect. The response of adult CD34^+^ HSPC controls to TGFβR1 inhibition was greater than RUNX1^in^ CD34^+^ HSPCs in colony assays (Figure 7B), suggesting additional TGFβR1-independent pathways may be involved. Nonetheless, these results support our hypothesis that elevated TGFβ1-associated and JNK2 signaling pathways in RUNX1^+/−^ contribute to defective megakaryopoiesis.

In summary, we demonstrate that RUNX1^+/−^ causes defects in early hematopoietic cells and in Mk lineage differentiation. RUNX1 activity in these cells is important for repression of TGFβR1/JNK2-associated signaling, and proinflammatory pathways that inhibit Mk progenitor differentiation and expansion. These studies, which extend RUNX1 function in terminal megakaryopoiesis to also promoting the earliest stages of Mk specification from HSPCs, have implications for developing better human models of FPDMM and provide insights that may lead to therapeutic strategies to alleviate the platelet defects and perhaps prevent the preleukemic evolution in FPDMM. Further work is needed to determine if blocking aberrant signaling in RUNX1^+/−^ progenitor cells will restore platelet counts and suppress progression of progenitor cells towards myelodysplastic/leukemic transformation in FPDMM patients.

## Authorship

BE carried out and evaluated these studies, prepared the first draft and subsequent revisions. SB constructed the L1, L1-C and WT-L1 lines, contributed to the methodology section and helped in useful discussions. DJ helped in early studies and their interpretation as well as manuscript editing. VS performed many of the western blot studies and their analysis. MG and PGao provided assistance in design and technical considerations of scRNA-SEQ experiments, MG, HH, JG assisted in data preparation and analyses, and PGao also assisted in preparation of scRNA-SEQ libraries. KT provided guidance in bioinformatic analysis. PL provided L2 and L2-C and contributed to editing of the manuscript. SBagga, NH, and WT provided assistance with technical considerations of generating lentiviral shRNA, as well as construction and testing of the NT- and RUNX1-targeting MSCV lentiviral vectors. NS provided guidance of the studies and assisted in editing of the manuscript. DF and PG guided the construction of lines L1, L1-C, WT, WT-Mut and helped provide overall guidance as well as manuscript editing. MP provided overall project organization and direction, data interpretation, and manuscript preparation. None of the authors have any financial or related disclosures to make with respect to this manuscript.

## Acknowledgments

This work was supported by a National Heart Lung and Blood Institute grant T32HL007150 (BE) and RUNX1 Research Program grants in an association with the Alex Lemonade Foundation (MP, DF, PG, WT and NS), and the Fred Hutchinson Cancer Research Center Cooperative Centers Excellence in Hematology’s NIDDK Grant U54 DK106829.

## References

1. Okuda T, van Deursen J, Hiebert SW, Grosveld G, Downing JR. AML1, the Target of Multiple Chromosomal Translocations in Human Leukemia, Is Essential for Normal Fetal Liver Hematopoiesis. Cell. 1996;84(2):321–330.

2. Wang Q, Stacy T, Binder M, Marin-Padilla M, Sharpe AH, Speck NA. Disruption of the Cbfa2 gene causes necrosis and hemorrhaging in the central nervous system and blocks definitive hematopoiesis. Proc Natl Acad of Sci U S A. 1996;93(8):3444–3449.

3. Ichikawa M, Asai T, Saito T, et al. AML-1 is required for megakaryocytic maturation and lymphocytic differentiation, but not for maintenance of hematopoietic stem cells in adult hematopoiesis. Nature Med. 2004;10(3):299–304.

4. Sun W, Downing JR. Haploinsufficiency of AML1 results in a decrease in the number of LTR-HSCs while simultaneously inducing an increase in more mature progenitors. Blood. 2004;104(12):3565–3572.

5. Schlegelberger B, Heller PG. RUNX1 deficiency (familial platelet disorder with predisposition to myeloid leukemia, FPDMM). Semin Hematol. 2017;54(2):75–80.

6. Godley LA. Inherited predisposition to acute myeloid leukemia. Semin in Hematol. 2014;51(4):306–321.

7. Gabbeta J, Yang X, Sun L, McLane MA, Niewiarowski S, Rao AK. Abnormal inside-out signal transduction-dependent activation of glycoprotein IIb-IIIa in a patient with impaired pleckstrin phosphorylation. Blood. 1996;87(4):1368.

8. Sun L, Mao G, Rao AK. Association of CBFA2 mutation with decreased platelet PKC-theta and impaired receptor-mediated activation of GPIIb-IIIa and pleckstrin phosphorylation: proteins regulated by CBFA2 play a role in GPIIb-IIIa activation. Blood. 2004;103(3):948–954.

9. Sun L, Gorospe JR, Hoffman EP, Rao AK. Decreased platelet expression of myosin regulatory light chain polypeptide (MYL9) and other genes with platelet dysfunction and CBFA2/RUNX1 mutation: insights from platelet expression profiling. J Thromb and Haemost. 2007;5(1):146–154.

10. Jalagadugula G, Mao G, Kaur G, Goldfinger LE, Dhanasekaran DN, Rao AK. Regulation of platelet myosin light chain (MYL9) by RUNX1: implications for thrombocytopenia and platelet dysfunction in RUNX1 haplodeficiency. Blood. 2010;116(26):6037–6045.

11. Bluteau D, Glembotsky AC, Raimbault A, et al. Dysmegakaryopoiesis of FPD/AML pedigrees with constitutional RUNX1 mutations is linked to myosin II deregulated expression. Blood. 2012;120(13):2708–2718.

12. Stockley J, Morgan NV, Bem D, et al. Enrichment of FLI1 and RUNX1 mutations in families with excessive bleeding and platelet dense granule secretion defects. Blood. 2013;122(25):4090–4093.

13. Glembotsky AC, Bluteau D, Espasandin YR, et al. Mechanisms underlying platelet function defect in a pedigree with familial platelet disorder with a predisposition to acute myelogenous leukemia: potential role for candidate RUNX1 targets. J Thromb and Haemost. 2014;12(5):761–772.

14. Latger-Cannard V, Philippe C, Bouquet A, et al. Haematological spectrum and genotype-phenotype correlations in nine unrelated families with RUNX1 mutations from the French network on inherited platelet disorders. Orphanet J Rare Dis. 2016;11:49–49.

15. Mao GF, Goldfinger LE, Fan DC, et al. Dysregulation of PLDN (pallidin) is a mechanism for platelet dense granule deficiency in RUNX1 haplodeficiency. J Thromb Haemost. 2017;15(4):792–801.

16. Rao AK, Poncz M. Defective acid hydrolase secretion in RUNX1 haplodeficiency: Evidence for a global platelet secretory defect. Haemophilia. 2017;23(5):784–792.

17. Jalagadugula G, Goldfinger LE, Mao G, Lambert MP, Rao AK. Defective RAB1B-related megakaryocytic ER-to-Golgi transport in RUNX1 haplodeficiency: impact on von Willebrand factor. Blood Adv. 2018;2(7):797–806.

18. Glembotsky AC, Sliwa D, Bluteau D, et al. Downregulation of TREM-like transcript-1 and collagen receptor alpha2 subunit, two novel RUNX1-targets, contributes to platelet dysfunction in familial platelet disorder with predisposition to acute myelogenous leukemia. Haematologica. 2019;104(6):1244–1255.

19. Woolthuis CM, Park CY. Hematopoietic stem/progenitor cell commitment to the megakaryocyte lineage. Blood. 2016;127(10):1242–1248.

20. Haas S, Trumpp A, Milsom MD. Causes and consequences of hematopoietic stem cell heterogeneity. Cell Stem Cell. 2018;22(5):627–638.

21. Noetzli LJ, French SL, Machlus KR. New insights into the differentiation of megakaryocytes from hematopoietic progenitors. Arterio Thromb Vasc Biol. 2019;39(7):1288–1300.

22. Sanjuan-Pla A, Macaulay IC, Jensen CT, et al. Platelet-biased stem cells reside at the apex of the haematopoietic stem-cell hierarchy. Nature. 2013;502(7470):232–236.

23. Yamamoto R, Morita Y, Ooehara J, et al. Clonal analysis unveils self-renewing lineage-restricted progenitors generated directly from hematopoietic stem cells. Cell. 2013;154(5):1112–1126.

24. Grover A, Sanjuan-Pla A, Thongjuea S, et al. Single-cell RNA sequencing reveals molecular and functional platelet bias of aged haematopoietic stem cells. Nature Comm. 2016;7:11075–11075.

25. Carrelha J, Meng Y, Kettyle LM, et al. Hierarchically related lineage-restricted fates of multipotent haematopoietic stem cells. Nature. 2018;554(7690):106–111.

26. Rodriguez-Fraticelli AE, Wolock SL, Weinreb CS, et al. Clonal analysis of lineage fate in native haematopoiesis. Nature. 2018;553(7687):212–216.

27. Miyawaki K, Iwasaki H, Jiromaru T, et al. Identification of unipotent megakaryocyte progenitors in human hematopoiesis. Blood. 2017;129(25):3332–3343.

28. Velten L, Haas SF, Raffel S, et al. Human haematopoietic stem cell lineage commitment is a continuous process. Nature Cell Biol. 2017;19(4):271–281.

29. Haas S, Hansson J, Klimmeck D, et al. Inflammation-Induced Emergency Megakaryopoiesis Driven by Hematopoietic Stem Cell-like Megakaryocyte Progenitors. Cell Stem Cell. 2015;17(4):422–434.

30. Tijssen MR, Cvejic A, Joshi A, et al. Genome-wide analysis of simultaneous GATA1/2, RUNX1, FLI1, and SCL binding in megakaryocytes identifies hematopoietic regulators. Dev Cell. 2011;20(5):597–609.

31. Beck D, Thoms JA, Perera D, et al. Genome-wide analysis of transcriptional regulators in human HSPCs reveals a densely interconnected network of coding and noncoding genes. Blood. 2013;122(14):e12–22.

32. Growney JD, Shigematsu H, Li Z, et al. Loss of Runx1 perturbs adult hematopoiesis and is associated with a myeloproliferative phenotype. Blood. 2005;106(2):494–504.

33. Soung do Y, Talebian L, Matheny CJ, et al. Runx1 dose-dependently regulates endochondral ossification during skeletal development and fracture healing. J Bone Miner Res. 2012;27(7):1585–1597.

34. Cai X, Gaudet JJ, Mangan JK, et al. Runx1 loss minimally impacts long-term hematopoietic stem cells. PloS one. 2011;6(12):e28430–e28430.

35. Connelly JP, Kwon EM, Gao Y, et al. Targeted correction of RUNX1 mutation in FPD patient-specific induced pluripotent stem cells rescues megakaryopoietic defects. Blood. 2014;124(12):1926–1930.

36. Antony-Debré I, Manchev VT, Balayn N, et al. Level of RUNX1 activity is critical for leukemic predisposition but not for thrombocytopenia. Blood. 2015;125(6):930–940.

37. Li Y, Jin C, Bai H, et al. Human NOTCH4 is a key target of RUNX1 in megakaryocytic differentiation. Blood. 2018;131(2):191–201.

38. Flohr T, Uppenkamp M, Baldus M, Hoffmann M, Huber C, Derigs HG. CD34+ cell enrichment for autologous peripheral blood stem cell transplantation by use of the CliniMACs device. J Hematotherapy Stem Cell Res. 2000;9(4):557–564.

39. Richel D, Johnsen H, Canon J, et al. Highly purified CD34+ cells isolated using magnetically activated cell selection provide rapid engraftment following high-dose chemotherapy in breast cancer patients. Bone Marrow Transplant. 2000;25(3):243.

40. Benjamini Y, Hochberg Y. Controlling the false discovery rate: a practical and powerful approach to multiple testing. Journal of the Royal statistical society: series B (Methodological). 1995;57(1):289–300.

41. Song WJ, Sullivan MG, Legare RD, et al. Haploinsufficiency of CBFA2 causes familial thrombocytopenia with propensity to develop acute myelogenous leukaemia. Nat Genet. 1999;23(2):166–175.

42. Mills JA, Paluru P, Weiss MJ, Gadue P, French DL. Hematopoietic differentiation of pluripotent stem cells in culture. Hematopoietic Stem Cell Protocols: Springer; 2014:181–194.

43. Paluru P, Hudock KM, Cheng X, et al. The negative impact of Wnt signaling on megakaryocyte and primitive erythroid progenitors derived from human embryonic stem cells. Stem Cell Res. 2014;12(2):441–451.

44. Ditadi A, Sturgeon CM, Keller G. A view of human haematopoietic development from the Petri dish. Nat Rev Mol Cell Biol. 2017;18(1):56–67.

45. Sullivan SK, Mills JA, Koukouritaki SB, et al. High-level transgene expression in induced pluripotent stem cell–derived megakaryocytes: correction of Glanzmann thrombasthenia. Blood. 2014;123(5):753–757.

46. Notta F, Zandi S, Takayama N, et al. Distinct routes of lineage development reshape the human blood hierarchy across ontogeny. Science. 2016;351(6269):aab2116.

47. Estevez B, Kim K, Delaney MK, et al. Signaling-mediated cooperativity between glycoprotein Ib-IX and protease-activated receptors in thrombin-induced platelet activation. Blood. 2016;127(5):626–636.

48. Estevez B, Du X. New concepts and mechanisms of platelet activation signaling. Physiology. 2017;32(2):162–177.

49. Becht E, McInnes L, Healy J, et al. Dimensionality reduction for visualizing single-cell data using UMAP. Nature Biotech. 2019;37(1):38–44.

50. Zhou Y, Zhou B, Pache L, et al. Metascape provides a biologist-oriented resource for the analysis of systems-level datasets. Nature Comm. 2019;10(1):1523.

51. Buettner F, Natarajan KN, Casale FP, et al. Computational analysis of cell-to-cell heterogeneity in single-cell RNA-sequencing data reveals hidden subpopulations of cells. Nature Biotech. 2015;33(2):155–160.

52. Lauridsen FKB, Jensen TL, Rapin N, et al. Differences in cell cycle status underlie transcriptional heterogeneity in the HSC compartment. Cell Reports. 2018;24(3):766–780.

53. Lu Y-C, Xavier-Ferrucio J, Wang L, et al. The molecular signature of megakaryocyte-erythroid progenitors reveals a role for the cell cycle in fate specification. Cell Reports. 2018;25(8):2083–2093. e2084.

54. Ahamed J, Janczak CA, Wittkowski KM, Coller BS. In vitro and in vivo evidence that thrombospondin-1 (TSP-1) contributes to stirring- and shear-dependent activation of platelet-derived TGF-beta1. PloS one. 2009;4(8):e6608–e6608.

55. Chen YZ, Incardona F, Legrand C, Momeux L, Caen J, Han ZC. Thrombospondin, a negative modulator of megakaryocytopoiesis. J Lab Clin Med. 1997;129(2):231–238.

56. Dave JM, Mirabella T, Weatherbee SD, Greif DM. Pericyte ALK5/TIMP3 axis contributes to endothelial morphogenesis in the developing brain. Dev Cell. 2018;44(6):665–678. e666.

57. Hofmann TG, Stollberg N, Schmitz ML, Will H. HIPK2 regulates transforming growth factor-β-induced c-Jun NH2-terminal kinase activation and apoptosis in human hepatoma cells. Cancer Res. 2003;63(23):8271–8277.

58. Miller MA, Sullivan RJ, Lauffenburger DA. Molecular pathways: receptor ectodomain shedding in treatment, resistance, and monitoring of cancer. Clinical Cancer Research. 2017;23(3):623–629.

59. Zhu L, Gomez-Duran A, Saretzki G, et al. The mitochondrial protein CHCHD2 primes the differentiation potential of human induced pluripotent stem cells to neuroectodermal lineages. J Cell Biol. 2016;215(2):187–202.

60. Botella LM, Sanz-Rodriguez F, Komi Y, et al. TGF-beta regulates the expression of transcription factor KLF6 and its splice variants and promotes co-operative transactivation of common target genes through a Smad3-Sp1-KLF6 interaction. Biochem J. 2009;419(2):485–495.

61. Zhu YX, Benn S, Li ZH, et al. The SH3-SAM adaptor HACS1 is up-regulated in B cell activation signaling cascades. J Exp Med. 2004;200(6):737–747.

62. Blank U, Karlsson S. TGF-β signaling in the control of hematopoietic stem cells. Blood. 2015;125(23):3542–3550.

63. Zhao M, Perry JM, Marshall H, et al. Megakaryocytes maintain homeostatic quiescence and promote post-injury regeneration of hematopoietic stem cells. Nature Med. 2014;20(11):1321–1326.

64. Kuter DJ, Gminski DM, Rosenberg RD. Transforming growth factor beta inhibits megakaryocyte growth and endomitosis.Blood. 1992 79(3):619–26.

65. van den Berk LC, Jansen BJ, Snowden S, et al. Cord blood mesenchymal stem cells suppress DC-T Cell proliferation via prostaglandin B2. Stem Cells Dev. 2014;23(14):1582–1593.

66. Zhang YE. Non-Smad signaling pathways of the TGF-β family. Cold Spring Harbor Perspec Biol. 2017;9(2):a022129.

67. Xiao X, Lai W, Xie H, et al. Targeting JNK pathway promotes human hematopoietic stem cell expansion. Cell discovery. 2019;5(1):2.

68. Angell RM, Atkinson FL, Brown MJ, et al. N-(3-Cyano-4,5,6,7-tetrahydro-1-benzothien-2-yl)amides as potent, selective, inhibitors of JNK2 and JNK3. Bioorg Med Chem Lett. 2007;17(5):1296–1301.

69. Huang N, Lou M, Liu H, Avila C, Ma Y. Identification of a potent small molecule capable of regulating polyploidization, megakaryocyte maturation, and platelet production. J Hematology Onco. 2016;9(1):136.

70. Team UoCHMCR. How I diagnose and manage individuals at risk for inherited myeloid malignancies. Blood. 2016;128(14):1800–1813.

71. Churpek JE, Bresnick EH. Transcription factor mutations as a cause of familial myeloid neoplasms. J Clin Invest. 2019;129(2):476–488.

72. Santini V, Valcárcel D, Platzbecker U, et al. Phase II Study of the ALK5 Inhibitor Galunisertib in Very Low-, Low-, and Intermediate-Risk Myelodysplastic Syndromes. Clin Cancer Res. 2019;25(23):6976–6985.

73. Sakurai M, Kunimoto H, Watanabe N, et al. Impaired hematopoietic differentiation of RUNX1-mutated induced pluripotent stem cells derived from FPD/AML patients. Leukemia. 2014;28(12):2344–2354.

74. Oved JH, Lambert MP, Kowalska MA, Poncz M, Karczewski K. Analysis of the Frequency of Spontaneous, Functionally-Significant Mutations in Genes Associated with Platelet Disorders in >120,000 Healthy Individuals. Blood. 2018;132(Supplement 1):2438–2438.

75. Gnatenko DV, Dunn JJ, McCorkle SR, Weissmann D, Perrotta PL, Bahou WF. Transcript profiling of human platelets using microarray and serial analysis of gene expression. Blood. 2003;101(6):2285–2293.

76. Shim MH, Hoover A, Blake N, Drachman JG, Reems JA. Gene expression profile of primary human CD34+CD38lo cells differentiating along the megakaryocyte lineage. Exp Hematol. 2004;32(7):638–648.

77. Hay SB, Ferchen K, Chetal K, Grimes HL, Salomonis N. The Human Cell Atlas bone marrow single-cell interactive web portal. Exp Hematol. 2018;68:51–61.

78. Psaila B, Wang G, Meira AR, et al. Single-cell analyses reveal aberrant pathways for megakaryocyte-biased hematopoiesis in myelofibrosis and identify mutant clone-specific targets. bioRxiv. 2019:642819.

