## Supplemental materials for "RUNX1 haploinsufficiency causes a marked deficiency of megakaryocyte-biased hematopoietic progenitor cells: Mechanistic studies and drug correction"

### Supplemental Methods

*Creating RUNX1$^{+/-}$ CD34$^{+}$-derived Mks*

Twenty-four hours prior to lentiviral transduction, adult, peripheral blood CD34$^{+}$ HSPCs were thawed and seeded at 3.3x10$^{5}$ per well in a 6-well plate coated with 40 µg/mL Retronectin (Takara Bio USA). Cells were seeded in fresh medium containing 80% Iscove's Modified Dulbecco's Medium (IMDM, Gibco containing 25mM HEPES, 25mM D-Glucose, 1mM Sodium Pyruvate) with GlutaMAX (4mM L-Alanyl-L-Glutamine), 20% BIT 9500 Serum Substitute and 100 mM βME supplemented with 300 ng/mL each of TPO, SCF and FLT3L (all, R&D Systems). After 24 hours, 10 multiplicity of infection (MOI) of virus solution containing *RUNX1*-silencing (clones V3SVHS02 _6883386 and V3SVHS02_10526718) or NT SMARTvector shRNA lentivirus (Dharmacon), each expressing GFP driven by the human elongation factor 1 alpha (*EF1A)* promoter (Figure S2) were added to cell cultures in combination with 1 µL LentiBlast transduction reagent (OZ Biosciences). After a 12-hour incubation, a second 10 MOI of the same viral stock was added. A different pair of *RUNX1*-targeting and non-targeting control shRNA were also generated and tested to confirm the SMARTvector RUNX1 shRNA. These lentiviral vectors are based on pCL20.MSCV.shLuc.PGK.mCherry[1,2] and were used to generate *RUNX1*-targeting or a NT luciferase control shRNA by cloning into the vector downstream of the murine stem cell virus (MSCV) promoter and upstream from a phosphoglycerate kinase (PGK) promoter driving mCherry (Figure S2). Cells were sorted for GFP or mCherry expression 72 hours after the first lentiviral transduction.

*Hematopoietic differentiation of iPSC- and CD34$^{+}$-derived progenitor cells*

For hematopoietic differentiation we used a previously established adherent feeder-free iPSC system[3,4]. Briefly, iPSCs were first depleted of mouse embryonic fibroblast feeder cells by treatment with 2mg/mL collagenase IV (Sigma-Aldrich) and passaging for 2 weeks. Feeder-depleted iPSCs were plated on Matrigel (BD Biosciences)-coated 6-well plates and cultured overnight with mTeSR (Stem Cell Technologies) media. The following morning (Day 0), media was exchanged with RPMI containing 1X Pen/Strep (Gibco, 100X), 2 mM glutamine (Gibco), 10 µg/mL ascorbic acid (Wako) and 4x10$^{-4}$ M monothioglycerol (MTG; Sigma-Aldrich) supplemented with BMP4 (5 ng/mL), VEGF (50 ng/mL) and 1 µM CHIR99021 for two days (Days 0-1). On day 2, media was replaced with RPMI containing BMP4 (5 ng/mL), VEGF (50 ng/mL) and bFGF (20 ng/mL) for two days (Days 2-3). On Day 4, media was replaced with SP34 (Gibco) containing 2mM Glutamine and Pen/Strep supplemented with BMP4 (5 ng/mL) , VEGF (15 ng/mL) and bFGF (5 ng/mL) for two days (Day 4-5). On Day 6, media was replaced with in homemade serum-free, hematopoietic differentiation medium (SFD)[3] containing 75% IMDM, 22% Ham's F-12 (Corning), 2mM Glutamine (Gibco), 0.05% bovine serum albumin (BSA, Sigma-Aldrich), 1% B-27 supplement without vitamin A (Gibco), and 0.5% N2 supplement (Gibco), and the following cytokines were added: VEGF (50 ng/mL), bFGF (100 ng/mL), TPO (50 ng/mL), Flt3L (25 ng/mL), SCF (25 ng/mL), and IL-6 (10 ng/mL).

Differentiated iPSCs yielded free-floating iHPCs on Day 7 of culture[3,4] . SFD media was used to replenish and collect iHPCs from Days 6-10. These iHPCs were seeded in methylcellulose methacult assays (Stem Cell Technologies) for an additional 10 days or in collagen-based megacult colony assays for 5 days (Stem Cell Technologies) and counted visually using a phase contrast microscope according to manufacturer recommendations. Megacult media contained IL-3, IL-6 and TPO. Mk colonies were identified as groups of 3-20 cells immunocytochemically stained pink (CD41 positive) and counterstained with Evan's Blue to show nuclei. In a subset of studies colonies with greater than 20 cells were scored as large Mk colonies. Methacult media contained SCF, GM-CSF, IL-3, IL-6 and G-CSF, and EPO. Colonies containing tightly packed clusters of 8 or more pale or dark reddish colored cells, due to hemoglobinization, were counted as erythroid colonies. Colonies containing clusters of 40 or more relatively loosely distributed cells appearing white-colored were counted as granulocyte/macrophage (GM) colonies.

For liquid cultures, Day 7 iHPCs were expanded for 8-9 days in homemade serum-free, hematopoietic differentiation medium (SFD)[3] containing contained 100 ng/mL TPO, 1 ng/mL SCF, 7.5 ng/mL interleukin 6 (IL-6) and 13.5 ng/mL IL-9 (all cytokines, R&D Systems). iMks were studied on Days 12 and 13 post-differentiation of iHPCs.

$CD34^+$ HSPCs were differentiated into Mks in 80% IMDM with GlutaMAX (Invitrogen), 20% BSA, recombinant human insulin (Stem Cell Technologies), and transferrin (BIT) 9500 Serum Substitute (Stem Cell Technologies) and 100 mM β-mercaptoethanol (βME, Sigma-Aldrich) containing 100 ng/mL TPO, 50 ng/mL SCF, and 7.5 ng/mL IL-6 and 13.5 ng/mL IL-9. On Day 10, SCF concentration was reduced to 1 ng/mL.[5] $CD34^+$-derived Mks were studied on Day 14.

To determine the effect on megakaryopoiesis from either the iPSC lines or $CD34^+$-derived HPCs, the following drugs were tested, each dissolved in DMSO: TGFβR1 inhibitors RS (100 nM)[6] and GS (100 nM)[7], γ-secretase indirect NOTCH inhibitor DAPT[8] (10 μM), and general JNK inhibitors, J-IN8 (100 nM, IN8)[9] and J-IX (300 nM)[10], all from Selleckchem.

*Fluorescent antibodies*

For surface staining and intracellular flowcytometry studies, specific combinations of antibodies conjugated to fluorescent molecules were used. Allophycocyanin-conjugated (APC) anti-CD235a , Brilliant ultraviolet 395 (BUV)-conjugated anti-CD41, eFluor450-conjugated anti-CD42a and phycoerythrin (PE)-conjugated anti-CD42b antibodies were used to isolate and enumerate iHPCs, whereas APC/Cyanine 7 (APC/Cy7)-conjugated anti-CD34, APC-conjugated anti-CD38,and PE conjugated anti-CD45RA, and PE/Dazzle 594-conjugated anti-CD41 antibodies were used to isolate and enumerate adult $CD34^+$ HSPCs. Absolute numbers of iPSC- or $CD34^+$-derived Mks were determined using fluorescent isothiocyanate (FITC)-conjugated annexin V in combination with CD41 and CD42 cell surface markers.

*scRNA-SEQ library preparation*

Day 7 L1-C and L1 iHPC samples were stained with antibodies against CD235a, CD41 and CD42a. Both were sorted into $CD42^-$ and $CD42^+$ iHPC fractions. Sorted cells were counted, and adjusted to a target of 5,000 cells in a 38 µL volume of 1X PBS (Gibco) and 0.04% BSA (Sigma-Aldrich). Cells, reverse transcription (RT) master mix (10X Genomics), primer-coated 10X gel beads, and partitioning oil were loaded onto a Single Cell A chip (10X Genomics) and into a 10X Genomics Chromium Controller instrument for microfluidic droplet-based encapsulation of cells, RT reagents, and gel beads. RT reaction was performed within droplets in a thermocycler (Bio-Rad) and following the 10X Genomics incubation protocol. Droplets were then broken down, and the released barcoded cDNAs were purified using magnetic beads, and amplified by PCR. cDNA concentrations were determine using a Qubit 3.0 fluorometer and Qubit HS dsDNA assay kit (both, ThermoFisher). Enzymatic digestion and size selection were performed to generate the optimal cDNA size for library construction. Sequencing primers P5, P7, R2 and sample-specific index primers were added during library construction. Quality and size of generated libraries was verified using the Bioanalyzer high-sensitivity DNA analysis kit (Agilent). Library concentrations were determined using kapa DNA library quantification kit (KAPA Biosystems). Libraries were sequenced on an Illumina HiSeq 4000 to obtain ~11,000 aligned reads per cell to identify cell types[11-13].

*scRNA-SEQ data pre-processing and analysis*

Demultiplexed FASTQ files were processed using the CellRanger count pipeline (10x Genomics, v.3.1.0[14]) for alignment of sequencing reads to the GRCh38 transcriptome and creation of feature-barcode matrices. The Seurat package (v.3.0.2[15]) within the R computing environment was used to aggregate and analyze the dataset. To account for possible batch effects, a canonical correlation analysis was used to integrate and aggregate the individual libraries. Gene expression matrices were log normalized and scaled, and a Wilcoxon rank sum test was used to determine differentially expressed genes between experimental groups. DE genes with a log2FC >0.59 or <-0.59 (>1.5 or <-1.5 FC) between L1-C and L1 iHPCs were considered significant when the P value was <0.05. Volcano plots of differential gene expression were generated in R using the Bioconductor (v.3.10) and Enhanced volcano (v.1.2.0[16]) packages. Mk-specific genes were further analyzed between the L1-C and L1 iHPC subpopulations. These selected genes were enumerated in Table S3 and compiled from published gene expression profiles of adult human Mk[17], Mk progenitor[18], and platelets[19], and included those known to be important in inherited bleeding disorders[20]. Functional and biological enrichment analyses were performed by uploading differentially expressed gene lists to the web-based Metascape analysis portal (http://metascape.org/gp/index.html), which integrates multiple gene set enrichment databases, including Kyoto Encyclopedia of genes and genomes, Reactome, and Gene Ontology[21]. Further validation of a subset of enrichment analyses was also performed using GSEA software (http://software.broadinstitute. org/gsea/index.jsp)[22].

Detailed gene expression analyses of the CD42a$^-$ and CD42a$^+$ L1-C or L1 iHPCs overall and within specific clusters are included in Figures S7 and S13, and Tables S7 and S8. Analyses of CD42$^-$ and CD42a$^+$ L1 iHPCs are in Figures S8 and S9, and Table S5).

*Western blotting of iMks*

Day 13 iMks were collected and sorted to enrich for CD42$^+$ iMks using anti-CD42 antibodies conjugated to phycoerythrin (PE), anti-PE-conjugated microbeads and MACS column for magnetic isolation of labeled cells (all, Miltenyi Biotec)[23]. Purified Mks were lysed with 1X radioimmunoprecipitation assay (1XRIPA) buffer containing 20 mM Tris-HCl (pH 7.5), 150 mM NaCl, 1 mM $Na_2$EDTA, 1 mM EGTA, 1% NP-40, 1% sodium deoxycholate, 2.5 mM sodium pyrophosphate, 1 mM β-glycerophosphate, 1 mM $NaVO_4$, and 1 mg/mL leupeptin (Cell Signaling Technology). The following inhibitors were also added to 1X RIPA: 5 mM phenylmethylsulfonyl fluoride (PMSF), 5 mM NaF, and 2.5 mM Halt phosphatase inhibitor (ThermoFisher). Final protein concentrations were determined using the Pierce BCA protein quantification kit (ThermoFisher). The sample protein (25 µg) was loaded into each well of a 10-well 4-12% precast Bis-Tris gradient gel (Invitrogen) as described[4] and run for 45-60 minutes at 135 mV. Sample proteins were then transferred to polyvinylidene fluoride (PVDF) membranes using an iBlot gel transfer system (Invitrogen). Subsequently, PVDF membranes were blocked in a 5% nonfat dry milk 1X Dulbecco's phosphate-buffered saline (1X DPBS, Gibco) solution (containing potassium chloride, potassium phosphate monobasic, sodium chloride, and sodium phosphate dibasic, without calcium or magnesium) for 30 minutes, rinsed briefly with 1X DPBS, and incubated in incubated in a 1:500 dilution containing one of the following: rabbit anti-phospho-SMAD 2/3, anti-SMAD 2/3, anti-JNK, anti-phospho-JNK, anti-P21, anti-P57, anti-TGFBR1 or 1:2000 anti-Vinculin or anti-GAPDH antibodies (all Cell Signaling Technologies, except anti-TGFBR1 from ThermoFisher) for 3 hours at room temperature. After primary antibodies were applied, membranes were washed 4 times for 5 minutes each in 1X DPBS, followed by incubation in horse radish peroxidase (HRP)-conjugated secondary antibodies for 1 hour in a 5% nonfat dry milk, 1X DPBS solution. Membranes were washed 4 times for 5 minutes each with a 1X DPBS + 0.5% Tween-20 solution after which, membranes were incubated with the SuperSignal West Femto enhanced chemiluminescent substrate (ThermoFisher), and light emitted from membrane-bound HRP-conjugated antibodies was visualized with X-ray film (LabScientific).

*Immunofluorescence detection of VWF, TLT-1 levels, and JNK and SMAD 2/3 phosphorylation*

Day 7 $CD42^-$ and $CD42^+$ iHPC sorted as described in *Flow Cytometry and Cell Sorting* above, were seeded in retronectin-coated 96 well plates for 2 hours, fixed and permeabilized with 1X BD Cytofix/Cytoperm reagent (BD Biosciences) according to the manufacturer recommendations. Cells were incubated with rabbit anti-VWF or Alexa Fluor 488 (AF488)-conjugated anti-TLT-1 antibodies at a 1:100 dilution overnight. In experiments where intracellular VWF and TLT-1 levels were studied by flow cytometry, Day 7 or Day 8 iHPCs were collected and stained with APC anti-CD235a , PE/Dazzle anti-CD41, and PE anti-CD42b (clone SZ2) antibodies for 15 minutes at room temperature prior to fixation and permeabilization.

When probing for intracellular phosphorylation, fixed/permeabilized samples were incubated overnight in a 1:50 dilution of rabbit anti-phospho-SMAD 2/3, anti-SMAD 2/3, anti-JNK or anti-phospho-JNK antibodies (see Table S1). After washing twice to remove excess primary antibodies with 1X Perm/Wash buffer (Diluted from a stock 10X Perm/Wash buffer containing fetal bovine serum and saponin, BD Biosciences), samples were incubated in the dark for 1 hour with goat anti-rabbit IgG AF488 (ThermoFisher). For intracellular flow cytometry, cells within the $CD42^-$ or $CD42^+$ $CD235^+CD41^+$ iHPC gate were analyzed for mean fluorescence intensity (MFI) of AF-488. When performing microscopy, 2 μg/mL Hoechst 33342 stain (ThermoFisher) was also added for the final 10 minutes of secondary antibody incubation. Samples were again washed twice and then stored at 4° C in 1X DPBS. Imaging was done using an EVOS FL Auto (Life Technologies) at 20X magnification.

### Supplement tables

**Table S1.** List of antibodies used in this paper. FC= flow cytometry. IF = immunofluorescence. WB = Western blot.

| Antigen | Conjugate | Clone | Vendor | Application |
|---|---|---|---|---|
| CD41 | APC | HIP8 | BD Pharmingen | FC |
| CD41 | BUV395 | HIP8 | BD Bioscience | FC |
| CD41 | PE/Dazzle-594 | HIP8 | Biolegend | FC |
| CD42b | PE | HIP1 | BD Pharmingen | FC |
| CD42b | PE | SZ2 | Beckman Coulter | FC/IF |
| CD42a | PE | ALMA.16 | BD Pharmingen | FC |
| CD42a | eFluor450 | GR-P | ThermoFisher | FC |
| CD235 | APC | HIR2 | BD bioscience | FC |
| VWF | None | Rb Polyclonal | Agilent/Dako | FC/IF |
| TLT-1 | AF488 | FAB 2394G | Novus Biologicals | FC |
| CD34 | APC/Cy7 | Gp105-120 | Biolegend | FC |
| CD38 | APC | HB7 | BD Biosciences | FC |
| CD45RA | PE | HI100 | BD Biosciences | FC |
| p-JNK | None | 81E11 | Cell Signaling | FC/WB |
| JNK | None | Rb Polyclonal | Cell Signaling | FC/WB |
| p-SMAD 2/3 | None | D27F4 | Cell Signaling | FC/WB |
| SMAD 2/3 | None | D7G7 | Cell Signaling | FC/WB |
| P21 | None | 12D1 | Cell Signaling | WB |
| P57 | None | Rb Polyclonal | Cell Signaling | WB |
| TGFβR1 | None | PA5-32631 | ThermoFisher | FC/WB |
| GAPDH | None | 14C10 | Cell Signaling | WB |
| Vinculin | None | E1E9V | Cell Signaling | WB |

**Table S2.** Total Mk yield from Day 7 iHPCs and percentage contribution of $CD42^+$ iHPC to total iMk yield

| | **Mk yield/day 7 $CD42^-$ iHPC (± 1 SEM)** | **Mk yield/day 7 $CD42^+$iHPC (± 1 SEM)** | **Mean frequency of day 7 $CD42^-$ iHPCs** | **Mean frequency of day 7 $CD42^+$ iHPCs** | **Yield of day 13 iMk/100 $CD42^-$ iHPCs** | **Yield of day 13 iMk/100 $CD42^+$ iHPCs** | **% Contribution of $CD42^+$iHPC to total day 13 iMK yield** |
|---|---|---|---|---|---|---|---|
| WT6 | 9 (± 2.9) | 6 (±1.9) | 88.7% | 11.3% | 834 | 72 | 8.7% |
| WT-L1 | 3 (± 0.6) | 2(± 0.8) | 98.5% | 1.51% | 286 | 4 | 1.4% |
| L1-C | 7 (± 1.5) | 9 (± 1.6) | 89.5% | 10.5% | 606 | 92 | 15% |
| L1 | 2 (± 0.3) | 4 (± 0.9) | 98.3% | 1.7% | 197 | 6 | 3% |

**Table S3.** See separate Excel file Table S3_RUNX1 for locally upregulated genes comparing specific $CD42a^+$ or $CD42a^-$ iHPC cell clusters in L1-C or L1.

**Table S4.** See separate Excel file Table S4_RUNX1 for manually curated 133 gene Mk/Plt gene list.

**Table S5.** See separate Excel file Table S5_RUNX1 for DE genes comparing all $CD42a^-$ versus $CD42a^+$ iHPCs in L1-C or L1.

**Table S6.** See separate Excel file Table S6_RUNX1 for lists of lineage selective and cell cycle genes to annotate cell clusters.

**Table S7.** See separate Excel file Table S7_RUNX1 for locally DE genes comparing L1-C and L1 within specific $CD42a^-$ or $CD42a^+$ iHPC cell clusters.

**Table S8.** See separate Excel file Table S8_RUNX1 for DE genes comparing all L1-C and L1 $CD42a^-$ or $CD42a^+$ iHPCs.

### Supplemental Figures

**A**

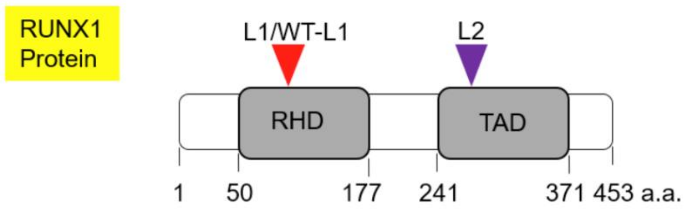


**B**

| iPSC line | *RUNX1* status | Affected protein domains |
|---|---|---|
| FPDMM L1 patient corrected (L1-C) | Corrected | None |
| FPDMM L1 patient (L1) | Exon 4, splice defect | RHD and TAD |
| CHOPWT6 (WT) | Normal | None |
| CHOPWT6 mutated (WT-L1) | Exon 4, splice defect | RHD and TAD |
| FPDMM L2 patient corrected (L2-C) | Corrected | None |
| FPDMM L2 patient (L2) | Exon 7 C→A transversion (Y260X) | TAD |

**Figure S1**. **Position of heterozygous *RUNX1* mutations in relation to RUNX1 protein structure and iPSC lines used in this study.**

(A) Schematic of RUNX1 protein structure highlighting the RUNT homology domain (RHD) and the transactivation domain (TAD)[24]. The location of the heterozygous frameshift present in lines L1 and WT-L1, and the nonsense RUNX1-truncating mutation in line L2 are noted. Amino acid numbering of isoform RUNX1b shown[25]. Heterozygous splice mutation present in L1 and WT-L1 results in loss of most of the RHD and everything downstream, including TAD. L2 retains the RHD, but truncates early in the TAD. All lines are predicted to result in $RUNX1^{+/-}$. (B) A table of all the $RUNX1^{+/-}$ mutant and paired control lines. Shown left to right is the name of the line and its abbreviation, the $\mathit{RUNX1}^{+/-}$ mutation and which domains are deleted. The shading designation used throughout the figures in the paper and supplement for the various lines is also indicated. The wildtype iPSC line WT was previously described as CHOPWT6[3,4]. FPDMM iPSC line 1 (L1) was derived from a patient previously described by us[26] and had a monoallelic splice acceptor defect near exon 4 of *RUNX*[26]. CRISPR/Cas9 technology[27] was used to monoallelically knockin the L1 mutation into the WT iPSCs (WT-L1) as well as to correct the mutation in L1 as we detailed (see manuscript Borst S, et al). The $RUNX1^{+/-}$ iPSC line L2 and its isogenic corrected control L2-C were previously described[28].

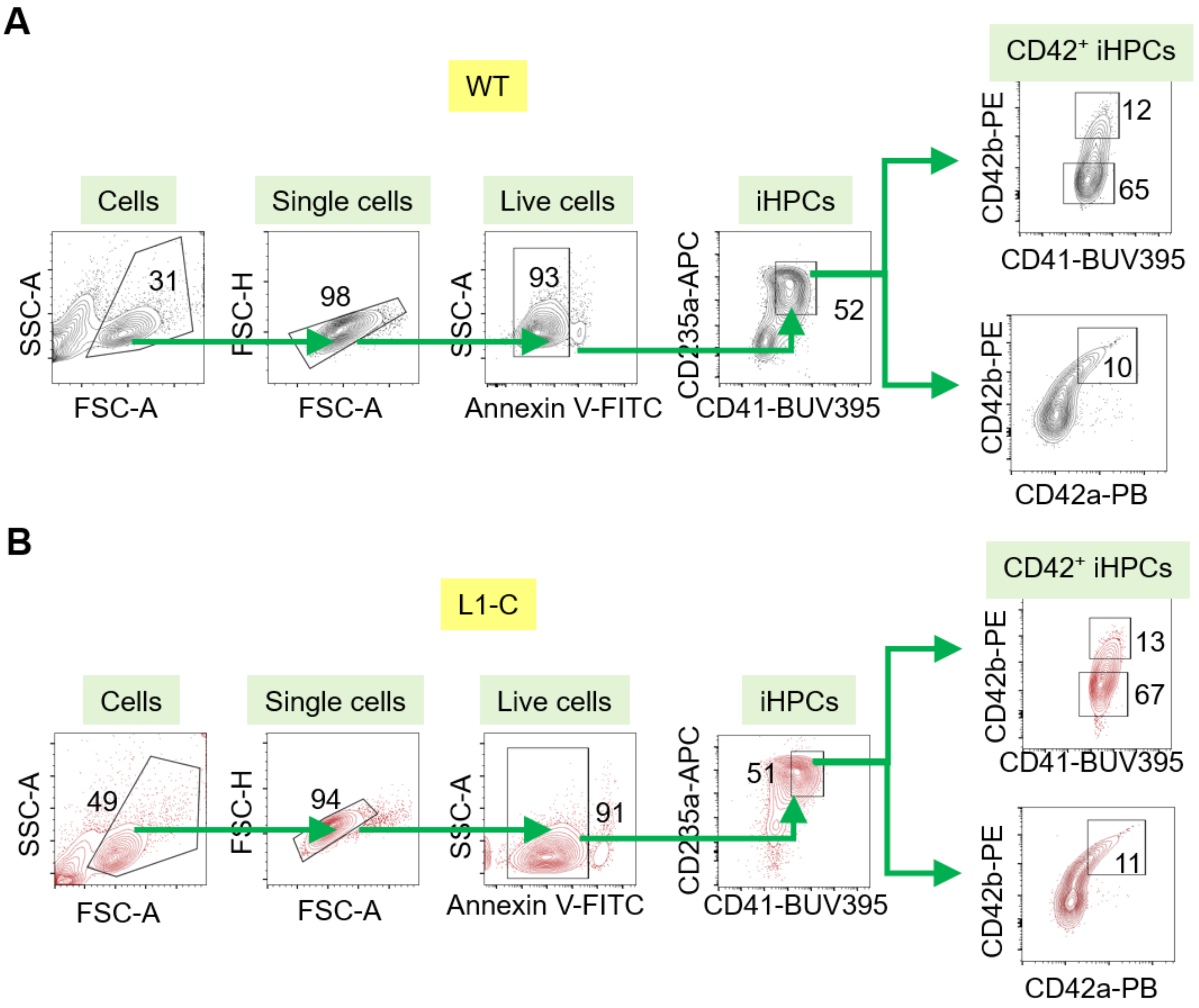


**Figure S2**. **Gating strategy for isolation and study of Mk-biased iHPCs.**
Representative flow cytometric analysis of Day 7 iHPCs stained for the indicated surface markers showing that ~10% of CD235$^{+}$/CD41$^{+}$ iHPCs express CD42 in a control (A) WT and (B) in a corrected FPD/AML line L1-C. Shown are detection of the GPIb/IX receptor with either anti-CD42a or CD42b antibodies.

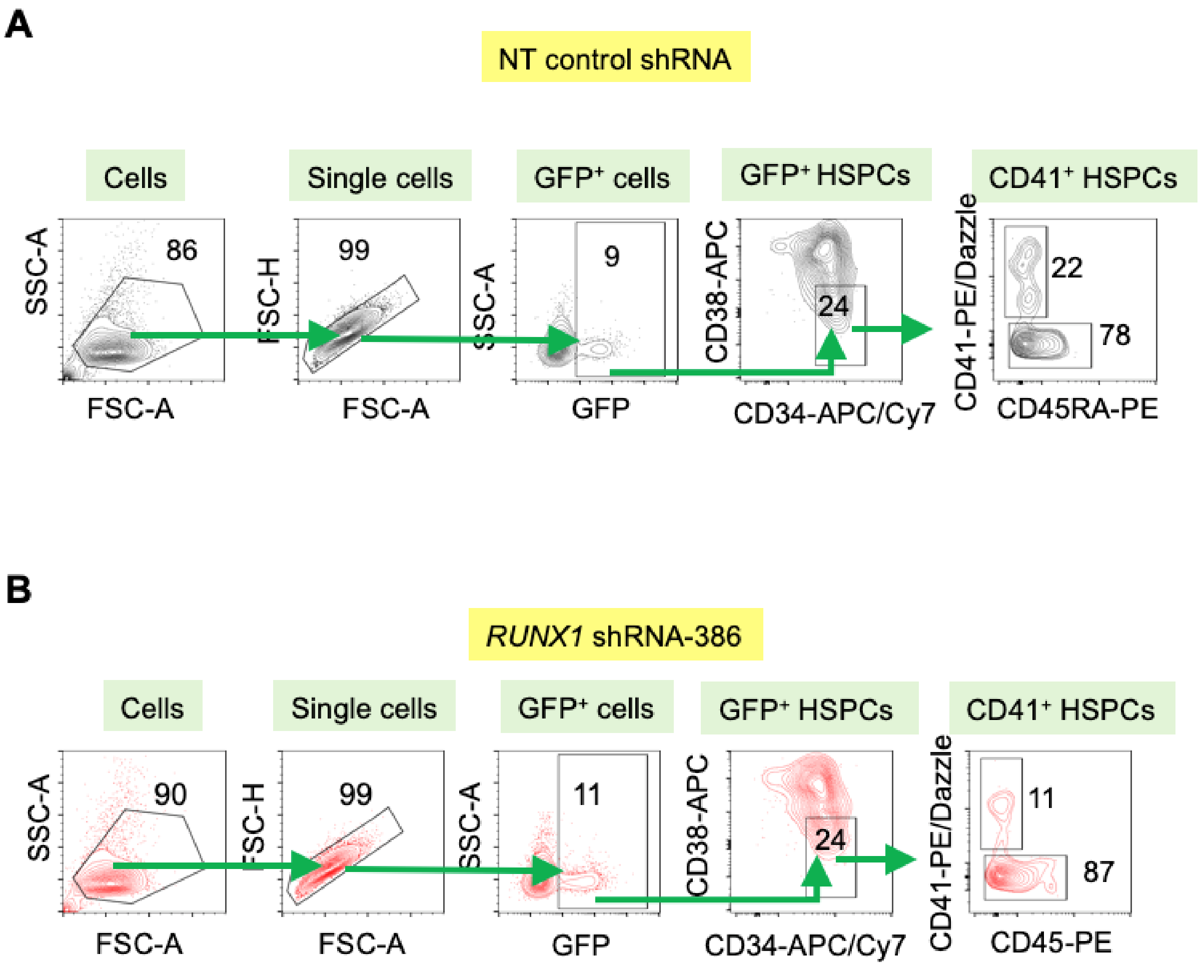


**Figure S3. Gating strategy for isolation and study of adult CD34$^+$ Mk-biased HPCs.**
Representative flow cytometric analysis of Day 4 cultured adult G-CSF-mobilized peripheral blood CD34$^+$ HSPCs stained for the indicated surface markers after transduction with lentiviruses expressing a GFP reporter. (A) NT shRNA-expressing cells showing 22% of control CD34$^+$CD38$^-$CD45RA$^-$ HSPCs express CD41 and a reduction in (B) *RUNX1* shRNA-expressing cells showing 11% of these HSPCs express CD41.

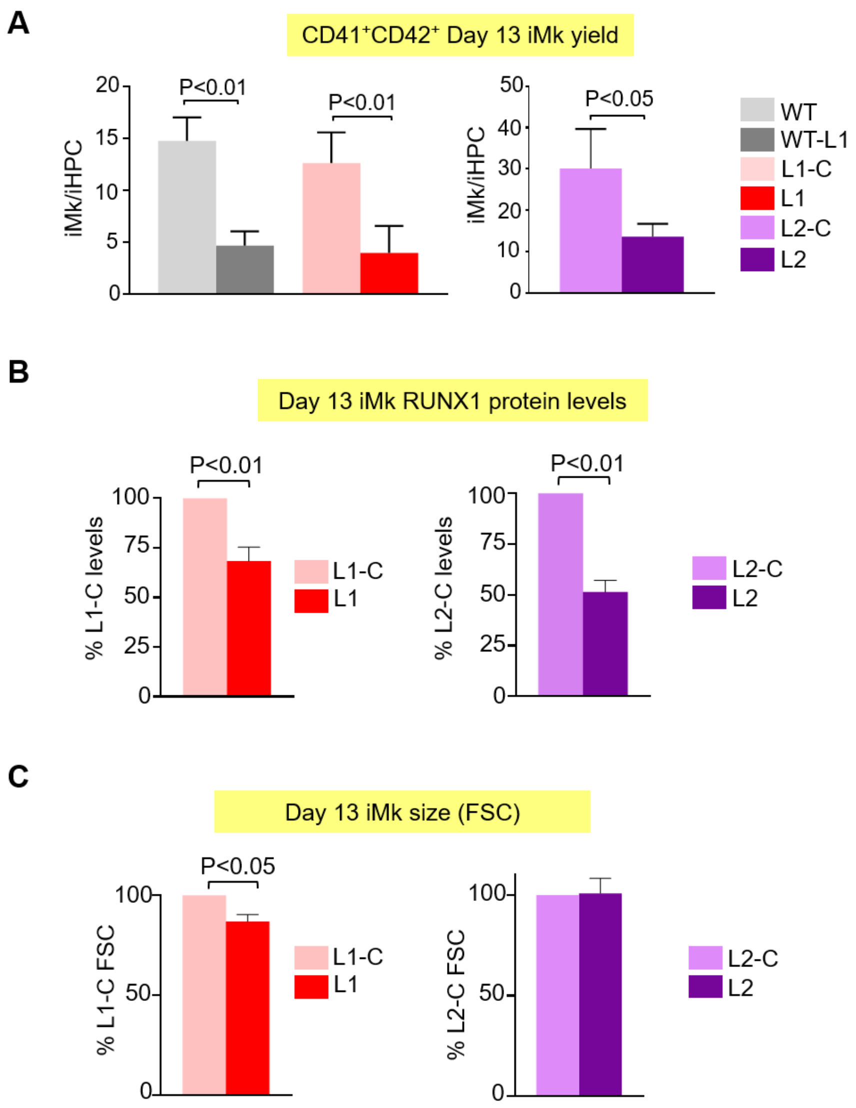


**Figure S4. RUNX1 protein levels and iMk yield from isogenic control and RUNX1$^{+/-}$ iPSC lines.** (A) Quantitation of the yield of Day 13 iMks normalized per input Day 7 iHPC for control and RUNX1$^{+/-}$ lines. Mean ± 1 SEM. N = 5-15 studies/arm. (B) Quantitation of intracellular RUNX1 levels in Day 13 iPSC-derived iMk cultures as determined by flow cytometry. Shown are percent of geometric MFI in control CD41$^+$CD42b$^+$ L1-C iMks set to 100%. (C) Quantitation of forward scatter (FSC) MFI in Day 13 iPSC-derived iMk cultures as determined by flow cytometry. Mean ± 1 SEM. With N = 3 studies/arm. P values in (A)-(C) were determined by Student's t-test.

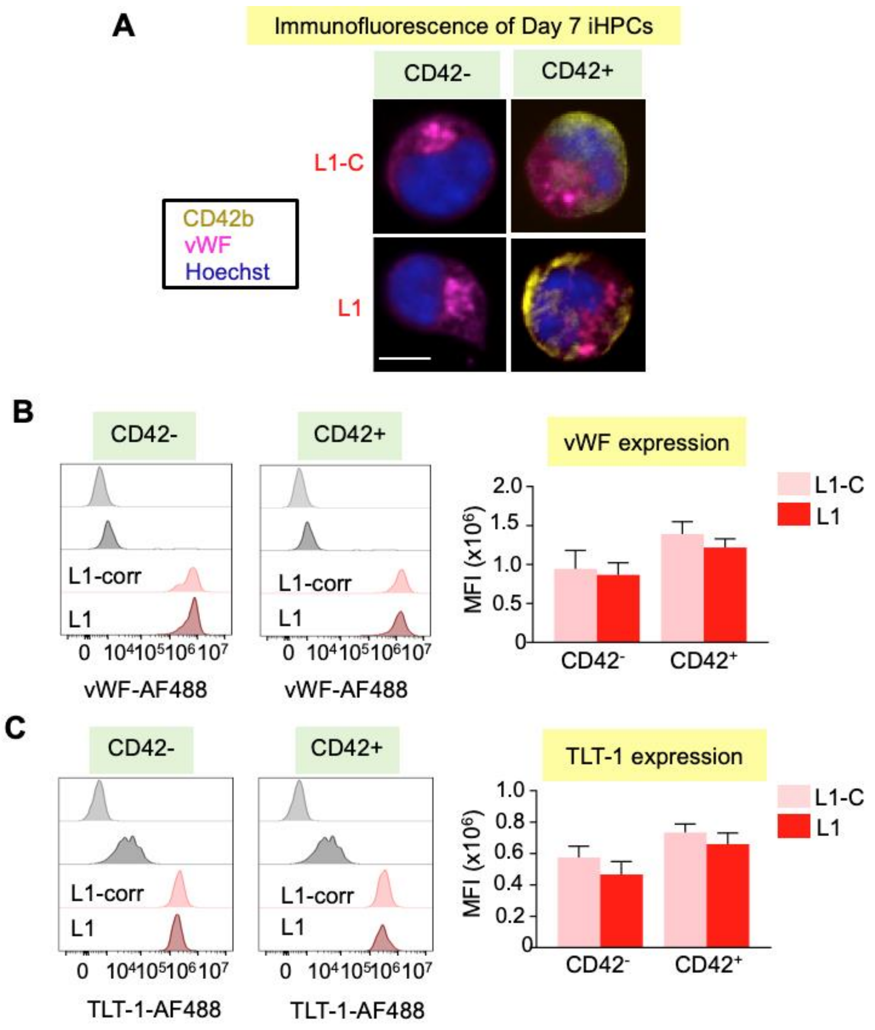


**Figure S5. Immunofluorescence and flow cytometric detection of VWF and TLT-1 in RUNX1$^{+/-}$ iHPCs.**

(A) Representative immunofluorescence microscopy of three studies sorted Day 7 CD42$^{-}$ and CD42$^{+}$ iHPCs from control L1-C and RUNX1$^{+/-}$ L1 lines. Sorted iHPCs were seeded in fibronectin-coated 96-well plates for 1 hour, stained with anti-CD42-PE antibodies (yellow), and then fixed and permeabilized. Cells were then stained with rabbit polyclonal anti-vWF antibodies, followed by anti-rabbit APC (magenta) and Hoechst (blue) and imaged on an EVOS FL microscope. Scale bar is 5µ. (B-C) Flow cytometric analysis of intracellular vWF and TLT-1 expression in sorted CD42$^{-}$ and CD42+ iHPCs. Left panels, overlayed fluorescence histograms for (B) vWF and (C) TLT-1 intracellular staining. Grey and black, denote unstained and secondary AF-488 antibody only control samples, respectively. Flow cytometry for intracellular staining was performed on Day 7 iHPCs after fixation and permeabilization, followed by staining with anti-VWF or anti-TLT-1 antibodies and then incubation overnight with anti-rabbit AF-488 secondary antibodies. Right panels, quantitation of MFI after iHPC immunostaining. Mean ± 1 SEM. N = 4 studies/arm analyzed by Student's t test.

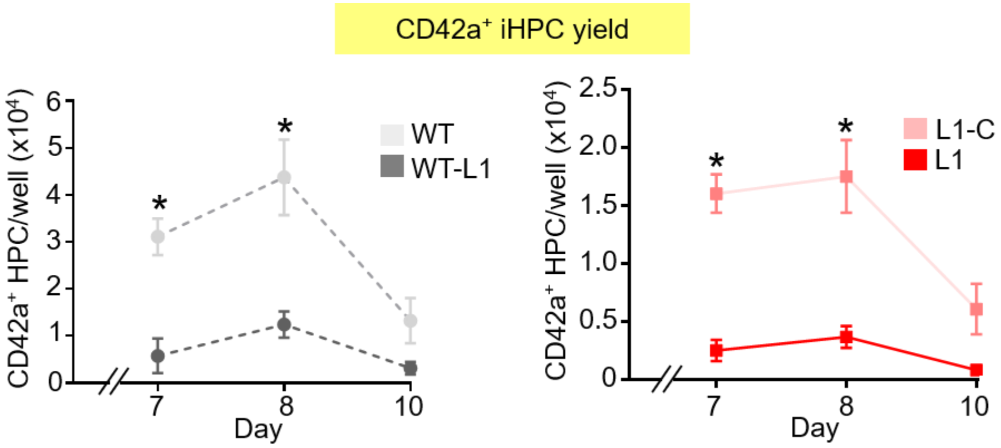


**Figure S6. Effect of RUNX1$^{+/-}$ on CD42a$^+$ iHPC size and kinetics.**

Isogenic control and RUNX1$^{+/-}$ iHPC studies. Mean yield of CD42a$^+$ iHPCs on Days 7, 8, and 10 of iPSC differentiation ± 1 SEM. N = 6 studies/arm. * = P<0.01 one-way ANOVA.

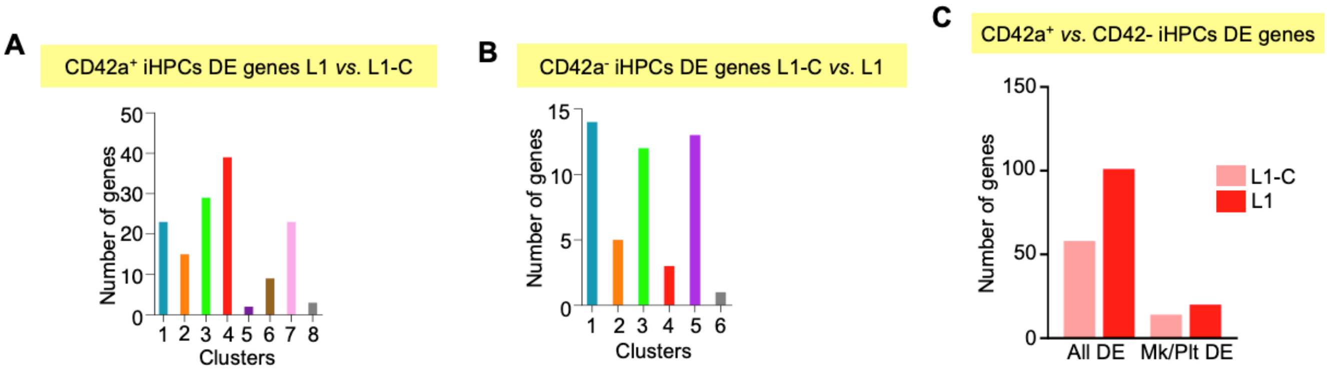


**Figure S7. Quantitation of DE genes per cluster in CD42a$^-$ and CD42a$^+$ iHPCs.**

Quantitation of the number of DE genes per cluster (A and B) or overall (C) across all cells in Day 7 iHPCs in L1-C and L1.

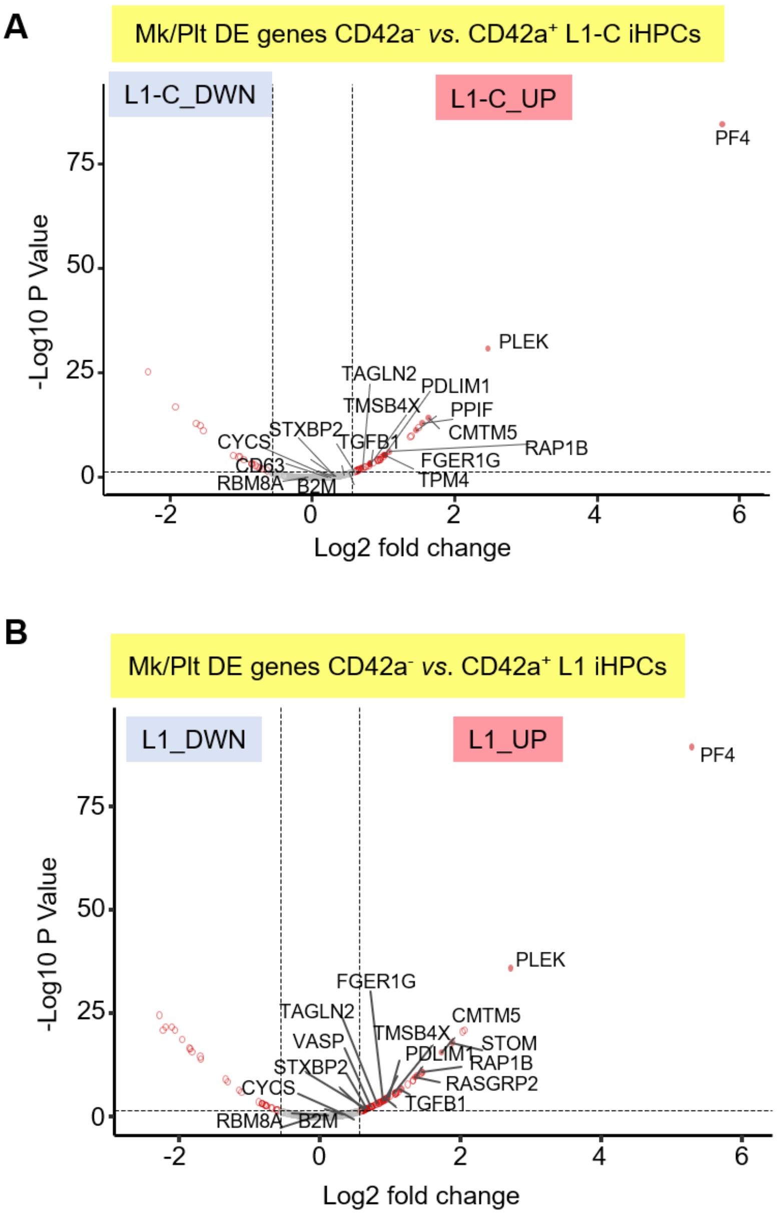


**Figure S8. Mk- and platelet-related genes enriched in L1-C and L1 CD42a$^+$ iHPCs.**

Volcano plots showing Mk/platelet (Plt) genes statistically enriched when comparing CD42a$^-$ to CD42a$^+$ iHPCs in (A) L1-C and (B) L1 iHPCs. Opened and red-filled-in circles are significant DE genes. Grey circles denote genes that did not meet statistical threshold for significance. Representative genes from a manually curated Mk/Plt list (Table S4) are filled in red. Note that only a subset of the Mk/Plt genes are noted. The full list of DE genes is in Table S5.

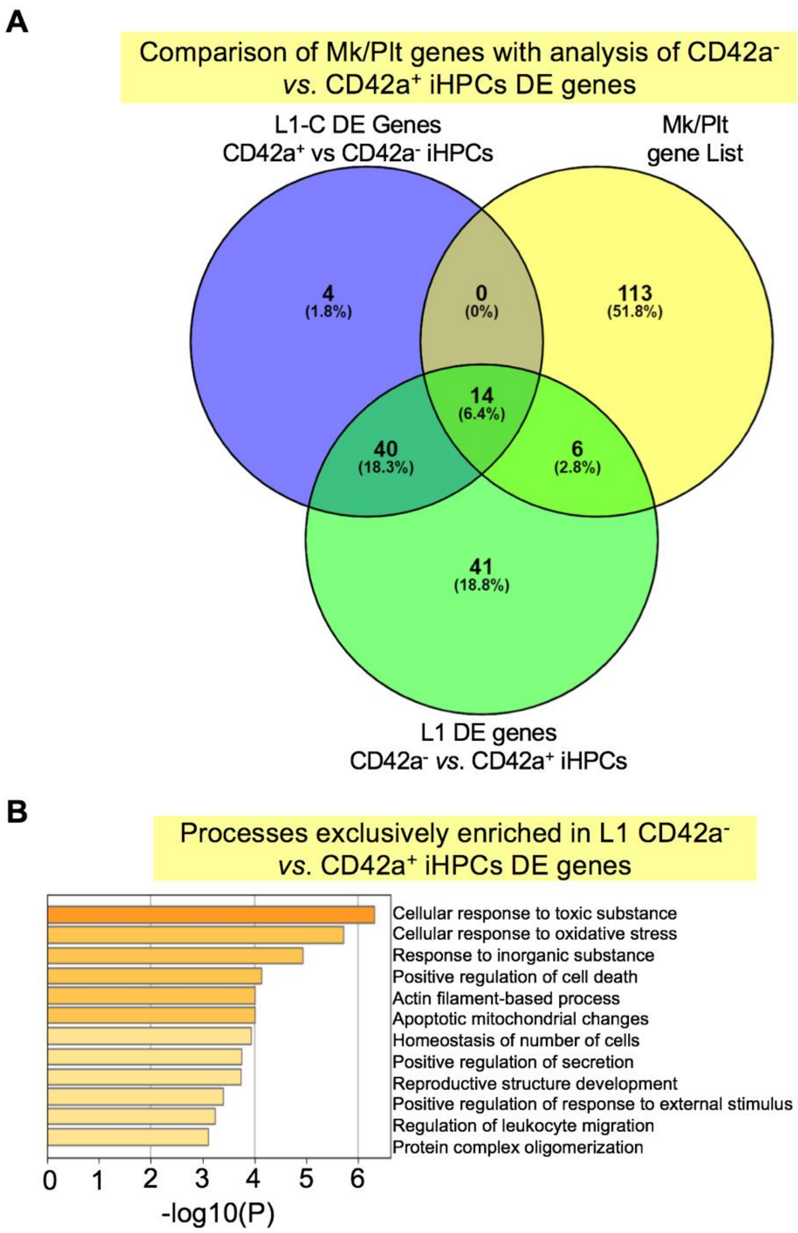


**Figure S9. Proportion of Mk/platelet genes enriched in CD42a$^+$ iHPCs and functional profile of DE genes exclusive to L1 CD42a$^+$ iHPCs**.

(A) Lists of DE genes in CD42a$^+$ versus CD42a$^-$ iHPCs from L1-C and L1 were used to generate Venn diagram and compared manually to a curated list of 133 Mk/platelet-associated genes. Note that most Mk/platelet genes are not DE in CD42a$^+$ iHPCs compared to CD42a$^-$ iHPCs in L1-C (14 genes) or L1 (20 genes). (B) Gene sets functionally enriched for stress response and immune-related processes in genes exclusively DE comparing CD42a$^+$ versus CD42a$^-$ L1 iHPCs. Similar enrichments are seen in direct DE comparison between L1 and L1-C shown in Figures 2C and 3C.

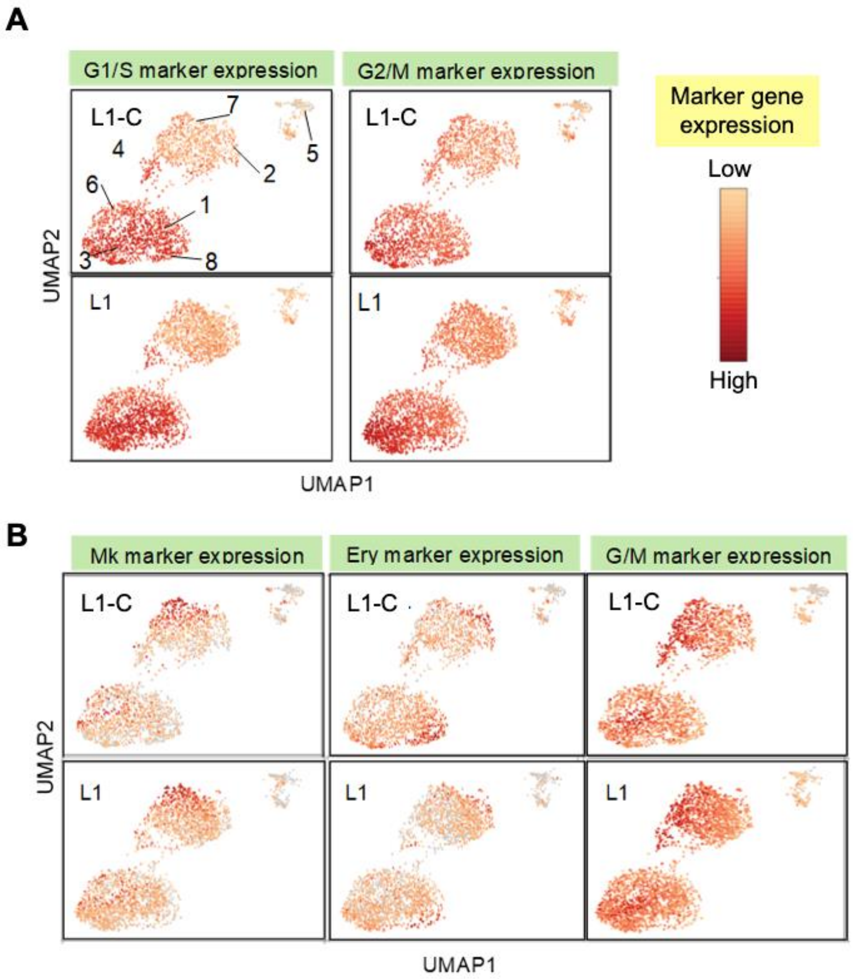


**Figure S10. Cell cycle and Mk, erythroid (Ery) and granulocyte/macrophage (G/M) marker gene expression in control LI-C and RUNX1$^{+/-}$ L1 CD42a$^{+}$ iHPCs.**

Expression of cell cycle and lineage maker gene sets was superimposed on the CD42a$^{+}$ iHPC UMAP plots. Distinct expression patterns of (A) cell cycle gene enrichment is seen in CD42a$^{+}$ iHPC 1, 3, 6 and 8 compared to (B) lineage marker enrichment in clusters 2, 4 and 7 which is indistinguishable between L1-C and L1 cells.

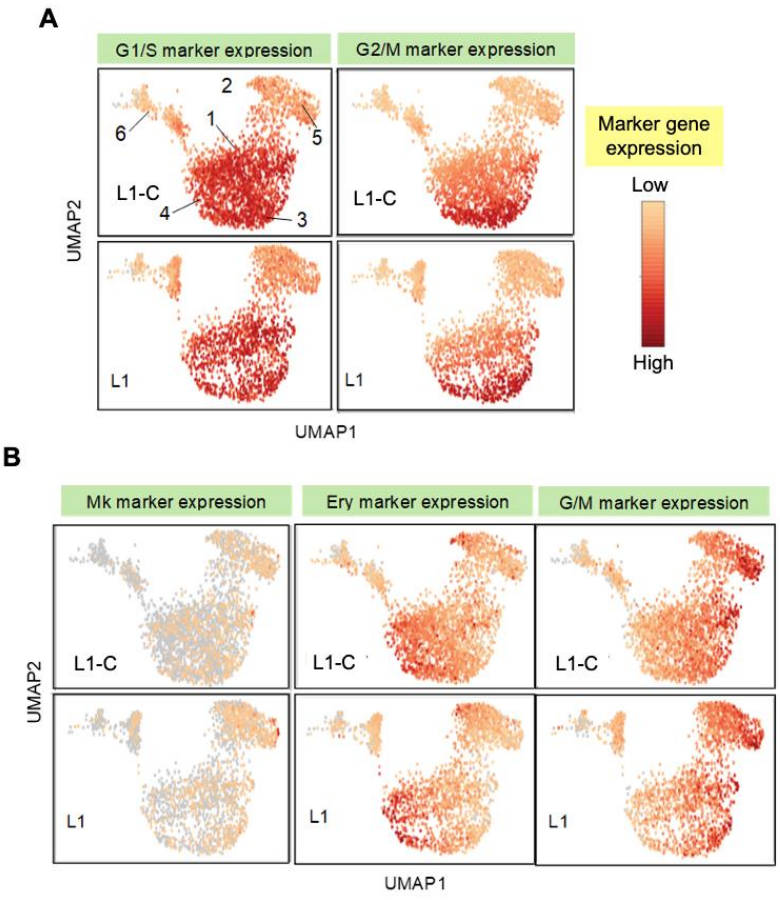


**Figure S11. Cell cycle and Mk, Ery and G/M marker gene expression in control LI-C and RUNX1$^{+/-}$ L1 CD42a$^{-}$ iHPCs.**

Expression of cell cycle and lineage maker gene sets was superimposed on the CD42a$^{-}$ iHPC UMAP plots. Distinct expression patterns of (A) cell cycle gene enrichment is seen in CD42a$^{-}$ iHPC clusters 1, 3, and 4 compared to (B) lineage marker enrichment in clusters 2 and 5 which is indistinguishable between L1-C and L1 cells.

A

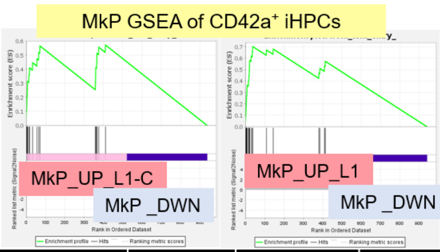


| MkP genes | NES | P-Value | FDR q-value |
|---|---|---|---|
| L1-C | 2.993 | 0 | 0 |
| L1 | 3.912 | 0 | 0 |

C

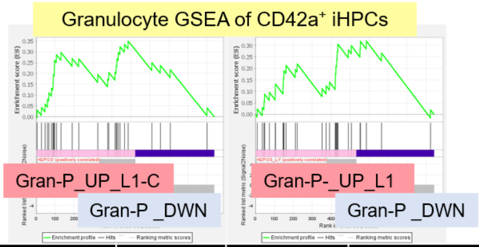


| Gran_genes | NES | P-Value | FDR q-value |
|---|---|---|---|
| L1-C | 2.184 | 0.004 | 0.004 |
| L1 | 1.956 | 0.008 | 0.008 |

B

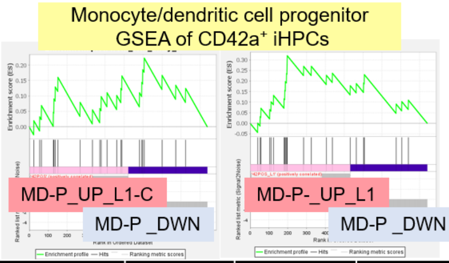


| MD-P genes | NES | P-Value | FDR q-value |
|---|---|---|---|
| L1-C | 1.235 | 0.208 | 0.208 |
| L1 | 1.803 | 0.014 | 0.014 |

D

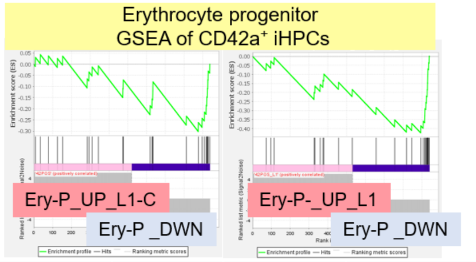


| EryP_genes | NES | P-Value | FDR q-value |
|---|---|---|---|
| L1-C | -1.766 | 0.004 | 0.014 |
| L1 | -2.547 | 0 | 0 |

**Figure S12. Gene set enrichment analysis of CD42a$^{+}$ iHPCs.**

Analysis by GSEA[22] of genes upregulated in CD42a$^{+}$ versus CD42a$^{-}$ iHPCs compared to a published adult human BM scRNA-SEQ data[18] containing genes sets associated with (A) MkPs, (B) MD-Ps, (C) granulocytes (Gran), and (D) EryP. The results show that control L1-C CD42a$^{+}$ iHPCs show strong positive enrichment for genes sets associated with MkP and granulocytes, weaker positive enrichment for MD-P genes, and negative correlation with EryP genes. NES = normalized enrichment score; FDR = false discovery rate.

**A**

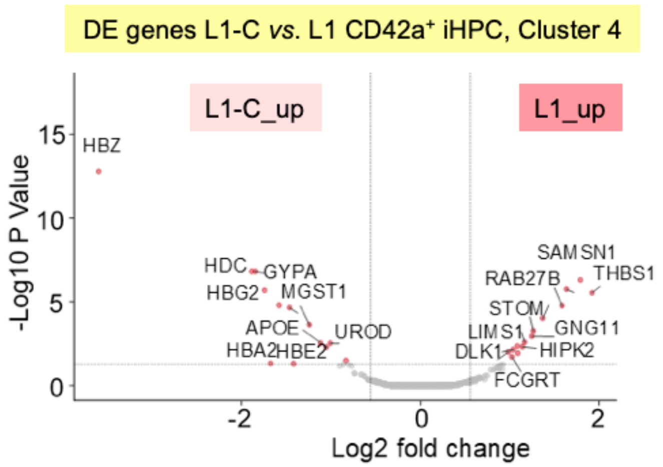


**B**

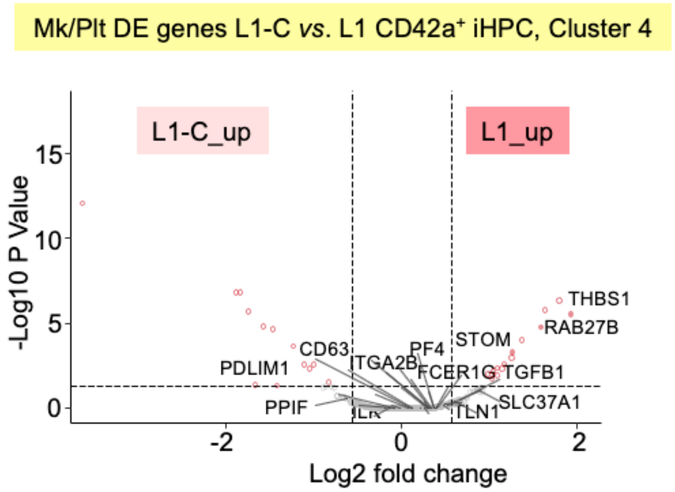


**Figure S13. DE gene analysis in L1 CD42a+ iHPC, Cluster 4.**

Volcano plots showing significantly changed genes (A) among all DE genes or (E) focusing only on Mk/Plt-associated DE genes when comparing L1-C to L1 CD42a+ iHPCs in Cluster 4. To improve clarity, not all genes are displayed in (A) only DE genes are shown, and in (B) most Mk/Plt genes are labeled (See Table S7 for full list DE genes in cell clusters). Note that most Mk/Plt genes are not significantly changed. Filled red circles denote significantly changed genes, whereas unfilled red circles indicate DE genes that are not on the list of Mk/Plt-associated genes.

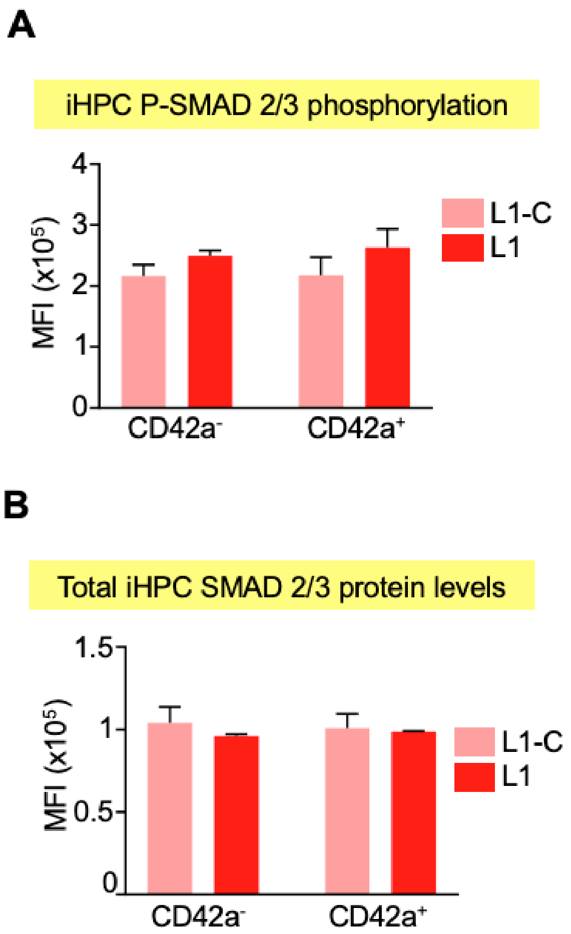


**Figure S14. Comparison of SMAD 2/3 activity and levels in L1-C and L1 iHPCs by intracellular flow cytometry.**

Quantitation of (A) SMAD 2/3 phosphorylation and (B) total SMAD 2/3 by intracellular flow cytometry. Day 8 iPSC-derived iHPCs were stained with iHPC surface markers, CD235 and CD41, and then with (A) anti-phospho-SMAD 2/3 or (B) anti-SMAD 2/3 (see Table S1). Cells within the $CD42^{+}$ or $CD42^{-}$ $CD235^{+}CD41^{+}$ iHPC gate were analyzed for MFI of AF-488. Shown are means ± 1 SEM with N = 3 studies/arm. Analysis was done by Student's t test.

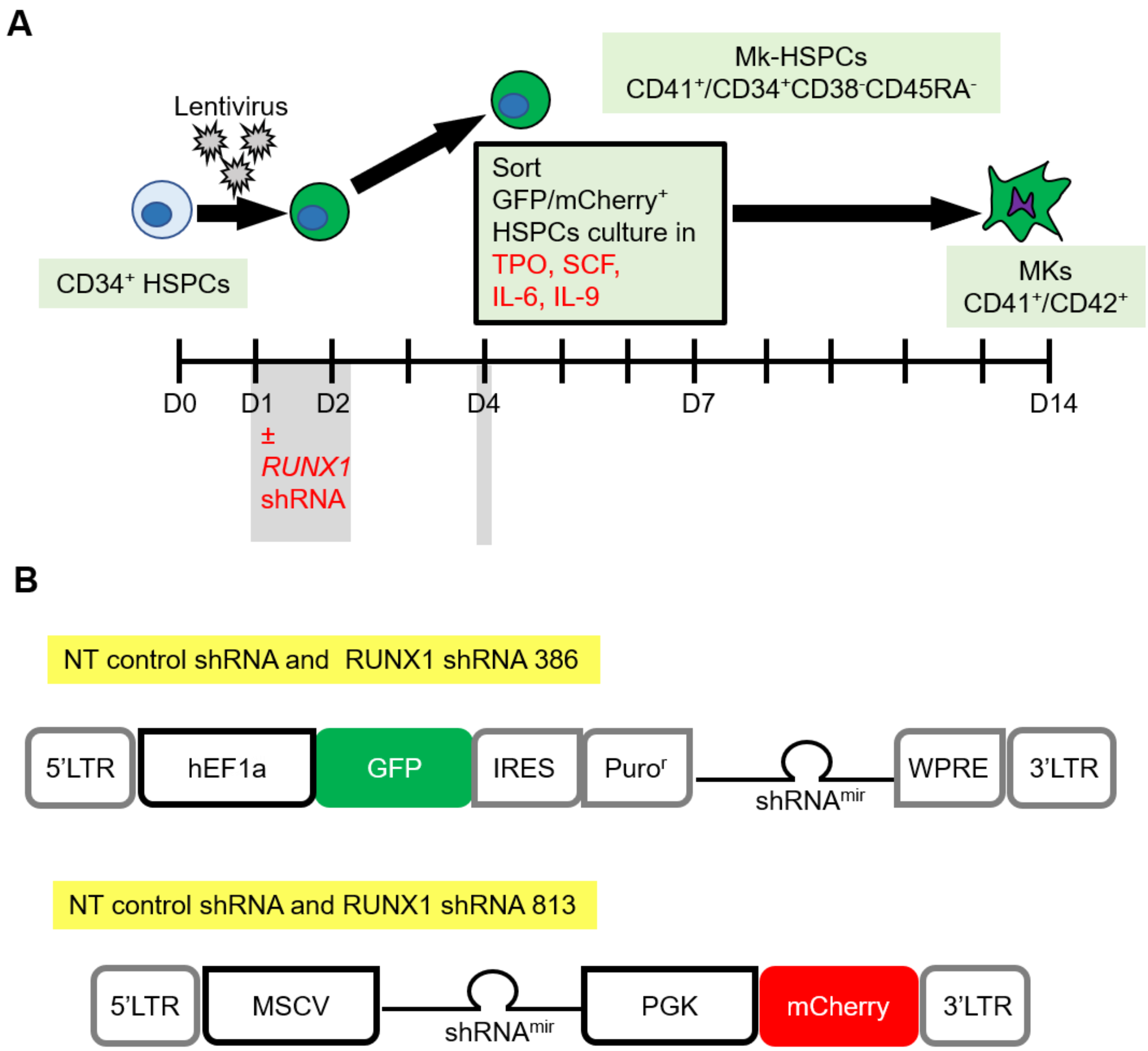


**Figure S15**. **Lentiviral transduction of adult CD34$^+$ HSPCs and shRNA constructs.**

(A) Schematic of how the shRNA lentiviral studies were done focused on Mk-biased HSPCs generated on Day 4 after transduction on Day 1. Mk-biased HSPCs were CD41$^+$/CD34$^+$CD38$^-$CD45RA$^-$. For studies on terminal Mks, cultures were harvested on Day 14. Timing of lentiviral transductions and cell sorting highlighted with grey shading. (B) *RUNX1* shRNAs were inserted into constructs downstream of a (top) human EF1a[1] or (bottom) MSCV promoter[1,2] in a microRNA (mir) scaffold. Elements of the SMARTvector (Dharmacon) or pCL20.MSCV.shLuc.PGK.mCherry vectors are shown. LTR = long terminal repeat; GFP = green fluorescent protein; IRES = internal ribosome entry site; Puro$^r$ = puromycin resistance; WPRE = woodchuck Hepatitis post-transcriptional regulatory element.

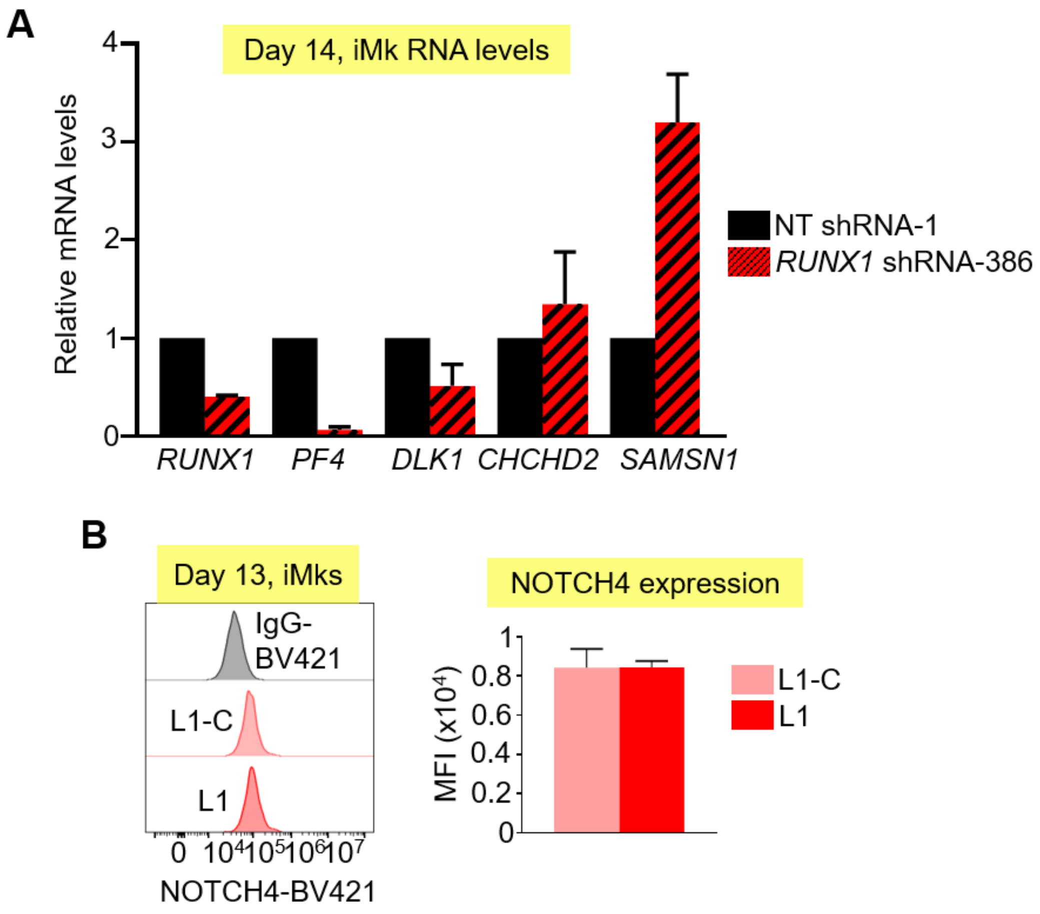


**Figure S16. Gene expression studies in Mks from *RUNX1*-targeting shRNA CD34$^{+}$ HSPCs and NOTCH4 levels in iMks from RUNX1$^{+/-}$ iHPCs.**

(A) Relative to NT shRNA transduction, the level in RUNX1$^{in}$ D14 Mks of *DLK1*, *CHCHD2* and *SAMSN1*. The latter two genes were differentially upregulated in RUNX1$^{in}$ cells by approximately 1.4 and 3-fold, respectively. *RUNX1* is shown demonstrating the efficacy of the shRNA approach and *PF4* is shown as a Mk-specific message. Mean ± 1 SEM are shown. N=2. (B) NOTCH4 levels in RUNX1$^{+/-}$ iMks. Left: Overlayed, representative, fluorescent histograms from flow cytometric analysis of NOTCH4 staining. Right: NOTCH 4 MFI by flow cytometry ± 1 SEM with N = 2 studies/arm.

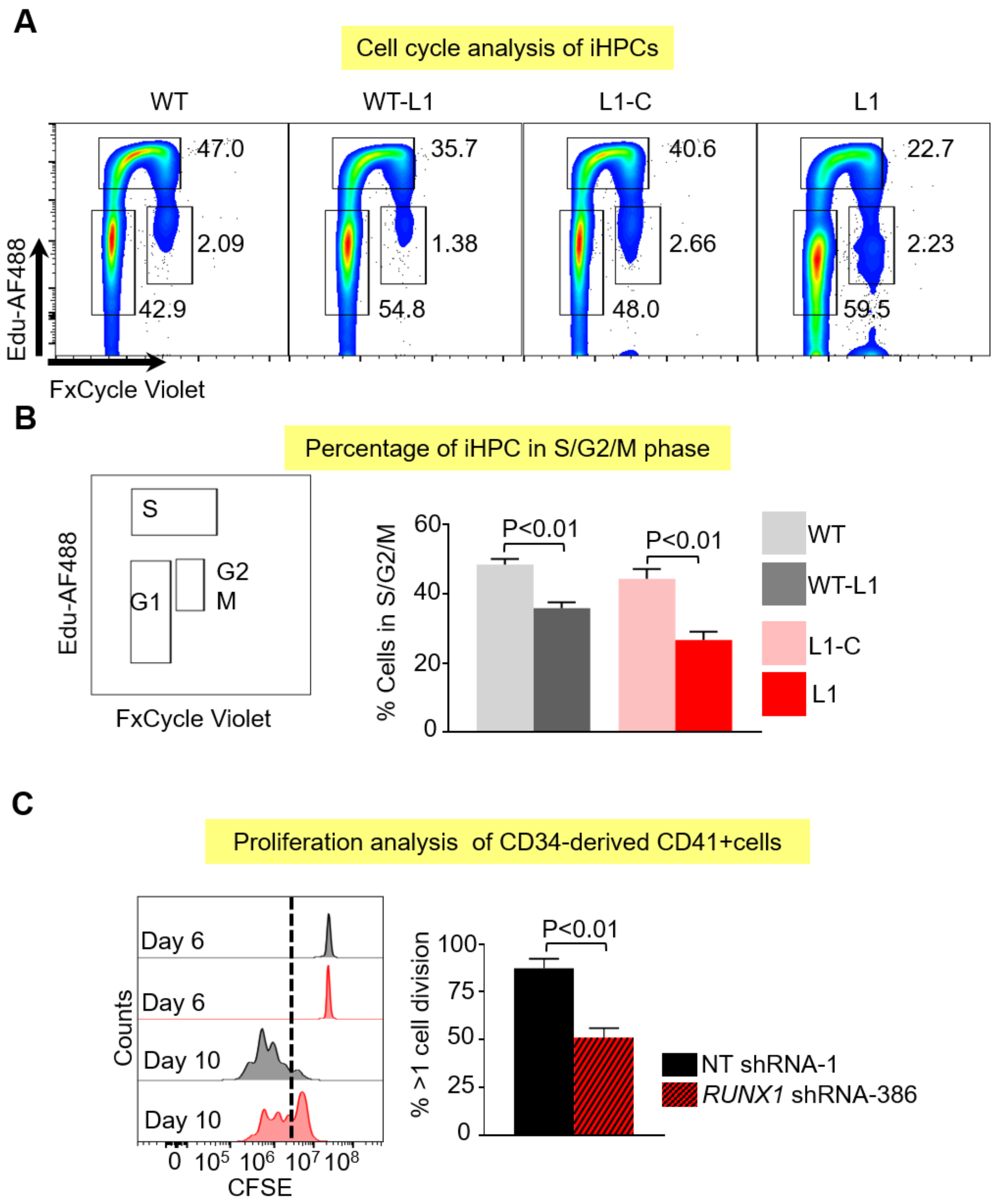


**Figure S17. Cell cycle studies in RUNX1$^{+/-}$ iHPCs and in RUNX1$^{in}$ adult CD34$^{+}$-derived cells.**

(A) Flow cytometric analysis of iHPCs 48hrs after thymidine analog Edu (5-ethynyl-2'-deoxyuridine) incorporation as detected by Click-it chemistry labeling with AF488 and co-staining with DNA-binding dye FxCycle-violet. (B) Left: Representative cell cycle gating strategy. Right: Quantitation of percentage of cells in S-phase and G2/M-phase. ± 1 SEM with N = 4 studies/arm. (C) Flow cytometric analysis of cell division in CD34-derived cells as determined by carboxyfluorescein diacetate, succinimidyl ester (CFSE) staining at baseline (Day 6) and its dilution four days later (Day 10). Left panels: Representative CFSE staining in

Day 6 and Day 10 CD34-derived CD41+ cells. Black, denotes non-targeting shRNA-1. Red, *RUNX1*-targeting shRNA-386. Dashed vertical line, shows cells having divided greater than or less than one time. Right panels, quantitation of percentage of cells dividing at least one time after CFSE staining. Mean ± 1 SEM. N = 4 studies/arm analyzed by Student's t test.

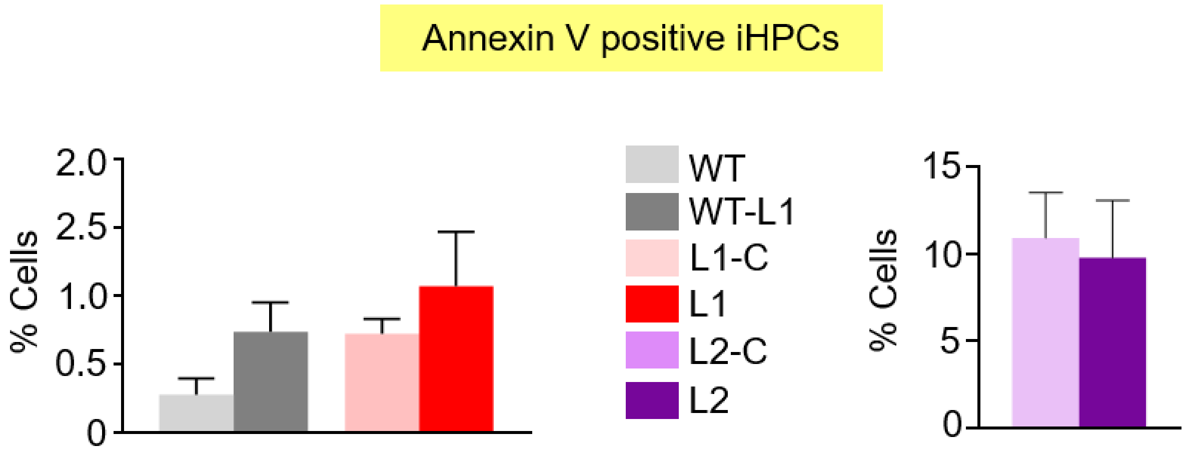


**Figure S18. Annexin V analysis in RUNX1$^{+/-}$ iHPCs.**

Flow cytometric analysis of apoptotic iHPCs. Day 7 iHPCs were collected and stained with Annexin V along with antibodies against iHPC markers CD41 and CD235a. Percentage of Annexin V positive iHPCs positive are shown. ± 1 SEM with N = 3-5 studies/arm.

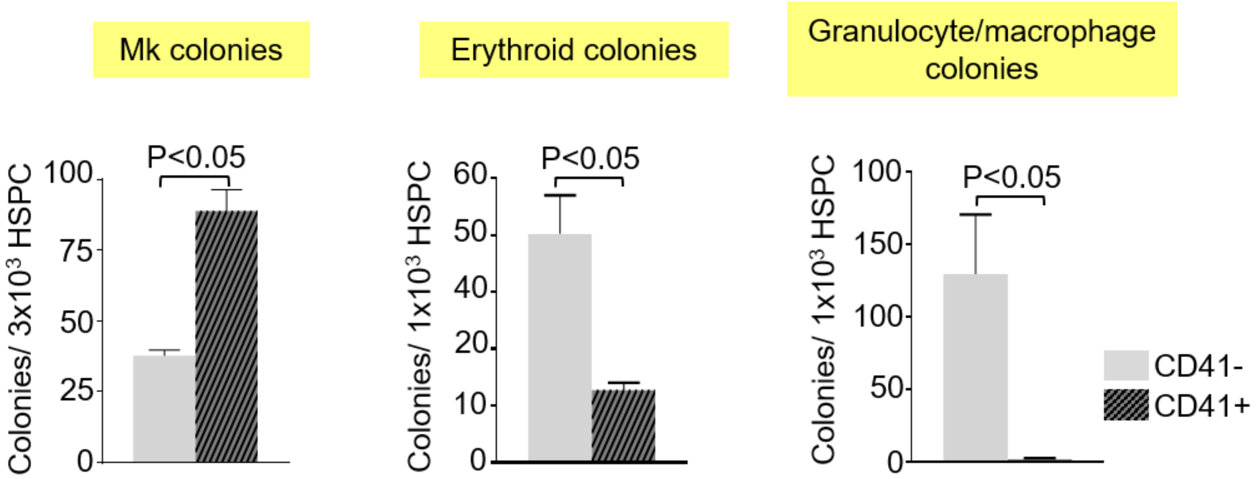


**Figure S19. Colony-forming assays using untransduced CD41$^-$ and CD41$^+$ HPSCs**

Untransduced CD41$^-$ and CD41$^+$ HPSCs were FAC-sorted and seeded in megacult media for Mk colonies or methacult media for erythroid colonies, and granulocyte/macrophage colonies. Mean ± 1 SEM are shown for N = 3 studies/arm analyzed by Student's t test.

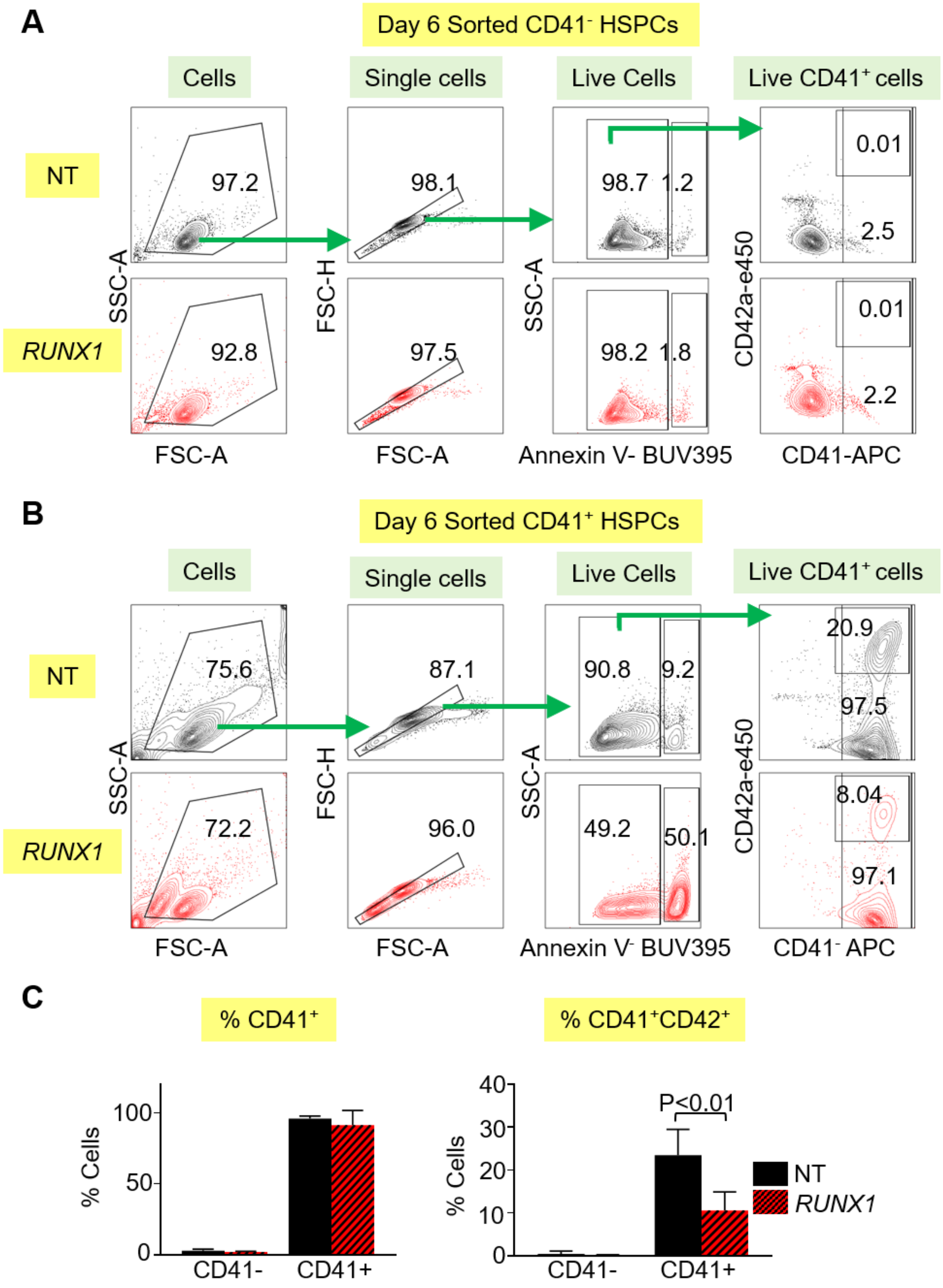


**Figure S20. Annexin V and Mk analysis in Day 6 sorted CD41- and CD41+ RUNX1$^{in}$ HSPCs.** (A-B) Gating strategy for flow cytometric analysis of sorted NT control shRNA-1 (NT) and *RUNX1*-targeting shRNA-386 (*RUNX1*) lentivirus expressing adult CD34-derived cells. Two days after cell sorting and culture in Mk media, on Day 6 (A) CD41- HSPCs, and (B) CD41+

HSPCs were stained for Annexin V, CD41 and CD42. (C) Left: Percentage of CD41+ cells and Right: Percentage of CD41+CD42+ cells on Day 6 of culture. Mean ± 1 SEM are shown for N = 3 studies/arm analyzed by Student's t test (related to Figure 6).

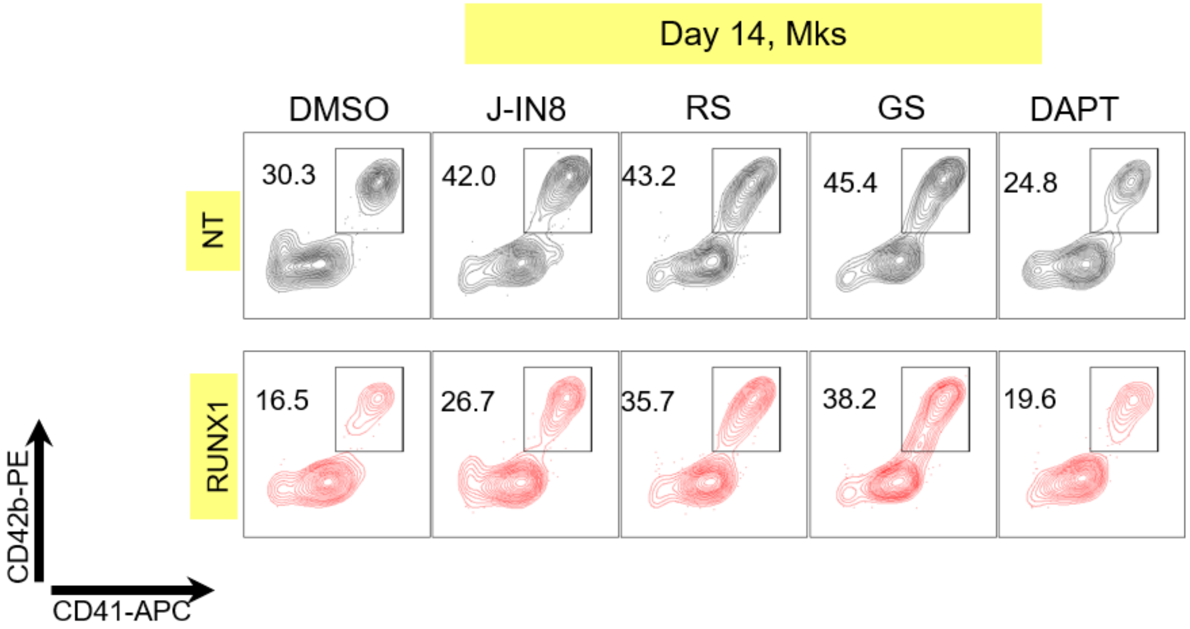


**Figure S21: Frequency of Day 14 Mks in drug-treated RUNX1$^{in}$ CD34-derived cells.** Representative flow cytometric analysis of drug-treated CD34-derived cells from NT shRNA control-1 and *RUNX1*-targeting shRNA-386 LV transduced HSPCs (related to Figure 7).

### References


1. Ramezani A, Hawley TS, Hawley RG. Lentiviral vectors for enhanced gene expression in human hematopoietic cells. *Mol Ther*. 2000;2(5):458-469.
2. Holmfeldt P, Ganuza M, Marathe H, et al. Functional screen identifies regulators of murine hematopoietic stem cell repopulation. *J Exp Med*. 2016;213(3):433-449.
3. Mills JA, Paluru P, Weiss MJ, Gadue P, French DL. Hematopoietic differentiation of pluripotent stem cells in culture. Hematopoietic Stem Cell Protocols: Springer; 2014:181-194.
4. Sullivan SK, Mills JA, Koukouritaki SB, et al. High-level transgene expression in induced pluripotent stem cell–derived megakaryocytes: correction of Glanzmann thrombasthenia. *Blood*. 2014;123(5):753-757.
5. Pineault N, Robert A, Cortin V, Boyer L. Ex vivo differentiation of cord blood stem cells into megakaryocytes and platelets. Basic Cell Culture Protocols: Springer; 2013:205-224.
6. Huang N, Lou M, Liu H, Avila C, Ma Y. Identification of a potent small molecule capable of regulating polyploidization, megakaryocyte maturation, and platelet production. *Journal of hematology & oncology*. 2016;9(1):136.
7. Akhurst RJ. Targeting TGF-beta Signaling for Therapeutic Gain. *Cold Spring Harb Perspect Biol*. 2017;9(10).
8. Olsauskas-Kuprys R, Zlobin A, Osipo C. Gamma secretase inhibitors of Notch signaling. *OncoTargets and therapy*. 2013;6:943-955.
9. Xiao X, Lai W, Xie H, et al. Targeting JNK pathway promotes human hematopoietic stem cell expansion. *Cell discovery*. 2019;5(1):2.
10. Angell RM, Atkinson FL, Brown MJ, et al. N-(3-Cyano-4,5,6,7-tetrahydro-1-benzothien-2-yl)amides as potent, selective, inhibitors of JNK2 and JNK3. *Bioorg Med Chem Lett*. 2007;17(5):1296-1301.
11. Streets AM, Huang Y. How deep is enough in single-cell RNA-seq? *Nat Biotechnol*. 2014;32(10):1005-1006.
12. Pollen AA, Nowakowski TJ, Shuga J, et al. Low-coverage single-cell mRNA sequencing reveals cellular heterogeneity and activated signaling pathways in developing cerebral cortex. *Nat Biotechnol*. 2014;32(10):1053-1058.
13. Jaitin DA, Kenigsberg E, Keren-Shaul H, et al. Massively parallel single-cell RNA-seq for marker-free decomposition of tissues into cell types. *Science*. 2014;343(6172):776-779.
14. Zheng GX, Terry JM, Belgrader P, et al. Massively parallel digital transcriptional profiling of single cells. *Nature communications*. 2017;8(1):1-12.
15. Butler A, Hoffman P, Smibert P, Papalexi E, Satija R. Integrating single-cell transcriptomic data across different conditions, technologies, and species. *Nature biotechnology*. 2018;36(5):411-420.
16. Blighe K. EnhancedVolcano: Publication-ready volcano plots with enhanced colouring and labeling. R package version 1.2. 0; 2019.
17. Shim MH, Hoover A, Blake N, Drachman JG, Reems JA. Gene expression profile of primary human CD34+CD38lo cells differentiating along the megakaryocyte lineage. *Exp Hematol*. 2004;32(7):638-648.
18. Hay SB, Ferchen K, Chetal K, Grimes HL, Salomonis N. The Human Cell Atlas bone marrow single-cell interactive web portal. *Exp Hematol*. 2018;68:51-61.
19. Gnatenko DV, Dunn JJ, McCorkle SR, Weissmann D, Perrotta PL, Bahou WF. Transcript profiling of human platelets using microarray and serial analysis of gene expression. *Blood*. 2003;101(6):2285-2293.
20. Oved JH, Lambert MP, Kowalska MA, Poncz M, Karczewski K. Analysis of the Frequency of Spontaneous, Functionally-Significant Mutations in Genes Associated with Platelet Disorders in >120,000 Healthy Individuals. *Blood*. 2018;132(Supplement 1):2438-2438.
21. Zhou Y, Zhou B, Pache L, et al. Metascape provides a biologist-oriented resource for the analysis of systems-level datasets. *Nature communications*. 2019;10(1):1523.

22. Subramanian A, Tamayo P, Mootha VK, et al. Gene set enrichment analysis: a knowledge-based approach for interpreting genome-wide expression profiles. *Proceedings of the National Academy of Sciences*. 2005;102(43):15545-15550.
23. Miltenyi S, Müller W, Weichel W, Radbruch A. High gradient magnetic cell separation with MACS. *Cytometry: The Journal of the International Society for Analytical Cytology*. 1990;11(2):231-238.
24. Kanno T, Kanno Y, Chen L-F, Ogawa E, Kim W-Y, Ito Y. Intrinsic transcriptional activation-inhibition domains of the polyomavirus enhancer binding protein 2/core binding factor α subunit revealed in the presence of the β subunit. *Molecular and cellular biology*. 1998;18(5):2444-2454.
25. Lam K, Zhang D-E. RUNX1 and RUNX1-ETO: roles in hematopoiesis and leukemogenesis. *Frontiers in bioscience: a journal and virtual library*. 2012;17:1120.
26. Song WJ, Sullivan MG, Legare RD, et al. Haploinsufficiency of CBFA2 causes familial thrombocytopenia with propensity to develop acute myelogenous leukaemia. *Nat Genet*. 1999;23(2):166-175.
27. Maguire JA, Cardenas-Diaz FL, Gadue P, French DL. Highly Efficient CRISPR-Cas9-Mediated Genome Editing in Human Pluripotent Stem Cells. *Current Protocols in Stem Cell Biology*. 2019;48(1):e64.
28. Connelly JP, Kwon EM, Gao Y, et al. Targeted correction of RUNX1 mutation in FPD patient-specific induced pluripotent stem cells rescues megakaryopoietic defects. *Blood*. 2014;124(12):1926-1930.
